# Growth and tension in explosive fruit

**DOI:** 10.1101/2023.05.15.540759

**Authors:** Gabriella Mosca, Ryan Eng, Milad Adibi, Saiko Yoshida, Brendan Lane, Leona Bergheim, Gaby Weber, Richard S. Smith, Angela Hay

**Affiliations:** Max Planck Institute for Plant Breeding Research, Carl-von-Linné-Weg 10, 50829 Köln, Germany; Technical University of Munich, 85748 Garching b. Munich, Germany; John Innes Centre, Norwich NR4 7UH, UK

**Author notes:** Present address: ZMBP, University of Tübingen, Auf der Morgenstelle 32, 72076 Tübingen, Germany. Present address: ETH, Otto-Stern-Weg 7, 8093/University of Zurich, Winterthurerstrasse 190, 8057 Zurich, Switzerland. Present address: University of Bern, Altenbergrain 21, 3013 Bern, Switzerland (Saiko Yoshida). Present address: HBRS, University of Applied Sciences, 53757 Sankt Augustin, Germany. These authors contributed equally to the work (Gabriella Mosca, Ryan Eng).

**Keywords:** Explosive seed dispersal, plant cell growth, cortical microtubules, cellulose microfibrils

## Abstract

Exploding seed pods of the common weed *Cardamine hirsuta* have the remarkable ability to launch seeds far from the plant. The energy for this explosion comes from tension that builds up in the fruit valves. Above a critical threshold, the fruit fractures along its dehiscence zone and the two valves coil explosively, ejecting the seeds. Tension is commonly generated as seed pods dry, causing fruit tissues to deform. However, this does not happen in *C. hirsuta.* Instead, tension is produced by active contraction of growing exocarp cells in the outer layer of the fruit valves. Exactly how growth leads to contraction in these cells is unknown. Here we show that microtubule dynamics in the exocarp cell cortex control the specific orientation of cellulose microfibrils in the cell wall, and the consequent cellular growth pattern, which together drive contraction. We used mechanical modeling and simulations to show how tension emerges through the general process of plant cell growth, due to the highly anisotropic orientation of load-bearing cellulose microfibrils and their effect on cell shape. By explicitly defining the cell wall as multi-layered in our model, we discovered that a cross-lamellate pattern of cellulose microfibrils further enhances the developing tension in growing cells. Therefore, the interplay of cell wall properties with turgor-driven growth enables the fruit exocarp layer to develop sufficient tension to explode.

## Introduction

Plants have evolved a multitude of different ways to disperse their seeds. Many of these dispersal structures have intriguing mechanical design features. For example, the pappus of the common dandelion has many closely spaced bristles, designed to generate a vortex air flow that optimizes the dispersal of seeds by wind [1]. Exploding seed pods, on the other hand, harness their own power to launch their seeds far from the plant. *Cardamine hirsuta* is related to the model plant *Arabidopsis thaliana*, but unlike *A. thaliana*, its seed pods explode. The elongated fruit of both species look very similar, comprising two valves that encase the seeds. In *C. hirsuta*, these valves coil rapidly outwards to launch the seeds at speeds greater than ten meters per second [2]. This rapid coiling relies on the differential contraction of valve tissues. Previous work showed that the outer exocarp cell layer of each valve actively contracts its reference length [2]. In contrast, the inner endocarp *b* cell layer of each valve is stiffened by a phenolic polymer called lignin that resists contraction [3]. Tension builds while the elongated valves are physically attached to the rest of the fruit. When structural elements of the fruit valves fail above a critical threshold, the valves rapidly coil to accommodate both the intrinsic contraction of the outer exocarp layer and the resistance to it from the inner lignified layer. It is crucial for the explosive mechanism that a substantial amount of tension develops in the exocarp cell layer, as this is linked to the velocity of seed launch.

Importantly, *C. hirsuta* fruit are characterized by their ability to explode without drying. This distinguishes them from many other explosive fruits, such as lupin, geranium and euphorbia, that generate tension through the differential shrinkage of drying tissues. The active contraction found in *C. hirsuta* fruit is a consequence of how exocarp cell shape deforms in response to the innate hydrostatic pressure found in plant cells called turgor [2]. Active osmoregulation alters turgor pressure and drives the swelling and shrinking of cells. This mechanism is used, for example in stomatal guard cells to open and close the stomatal pore, or in the specialized joint structures of Mimosa leaves to actuate leaf folding [4, 5]. However, the active contraction of *C. hirsuta* fruit exocarp cells is not associated with a change in turgor pressure [2], leaving open the question of how tension develops in the elongated fruit valves.

Turgor pressure is the major driving force behind expansive cell growth in plants. Plant cells are encased by a rigid yet malleable cell wall that functions to counter the large amount of turgor pressure found within a cell. This mechanical equilibrium can be turned into expansive growth if the cell wall is stretched beyond a certain threshold. This stretch induced by turgor, combined with active loosening of the cell wall [6], leads to irreversible expansion. Specifically, the relaxation of cell wall stress by loosening generates a slight reduction in water potential that allows cells to take up water and expand [7]. Mechanical equilibrium is re-established by new wall synthesis, and this continuous loop is reiterated, resulting in expansive growth [8, 9]. In this way, the precise interplay between turgor pressure and cell wall properties determines the growth of plant cells.

The main load-bearing components of the cell wall are cellulose microfibrils, each made up of many β- (1→4)-D-glucan polysaccharide chains, typically packed into a crystalline array. When cellulose microfibrils are aligned they bind tightly to each other and act as biological steel cables that resist being stretched [9]. In this way, cellulose microfibril alignment influences the direction of cellular growth. Since growth is restricted in the direction of aligned microfibrils, cells preferentially grow in the orthogonal direction. Cellulose microfibrils are synthesized by a heteromeric enzymatic complex composed of cellulose synthase subunits (CESAs) that are localized at the plasma membrane [10]. During cellulose biosynthesis, CESAs extrude microfibrils into the cell wall while moving within the plasma membrane. CESA motility is linear and is typically guided by the cortical microtubule cytoskeleton [11, 12], resulting in cellulose microfibrils that mirror microtubule orientation. Although there is evidence that CESA motility and cellulose microfibril synthesis may also be independent of microtubules [13, 14], it is widely accepted that microtubules determine cellulose microfibril alignment, which can then influence the direction of cellular growth and consequently cell shape [15, 16].

Here, we discovered that the fruit exocarp actively contracts in length and generates pulling force through normal cell growth. Since growth is a process that relaxes stress in the cell wall, it was unclear how growth could generate contractile tension. To address this problem, we used a combination of computational modeling, live confocal imaging, and microtubule perturbations. We found that a developmental switch in microtubule orientation is critical to reorient cellulose microfibrils and change cellular growth patterns in the fruit exocarp. Modeling indicates that active contraction emerges from the interaction of this oriented cellulose microfibril reinforcement and growth. By designing a multi-layered modeling framework, we found that a cross-lamellate pattern of cellulose microfibrils can further enhance the active contraction of growing exocarp cells.

## Results

### Cell shape, anisotropy and turgor-driven contraction

The elongated seed pods of *A. thaliana* and *C. hirsuta* look very similar despite their striking difference in seed dispersal (Fig. 1A). Yet, underlying these similar organ shapes are very different cell shapes. Exocarp cells, which comprise the epidermal layer of the fruit valve, have an elongated shape in *A. thaliana*, described by high aspect ratios with considerable variation (Fig. 1B, D). These same cells are uniformly square in *C. hirsuta* (Fig. 1B, D). Turgor pressure exerts mechanical stress on the cell wall and the extent of this stress depends on the shape and size of the cell. For example, large cells bulge out and experience more stress than small cells [17]. Exocarp cells in both *A. thaliana* and *C. hirsuta* grow larger by cell expansion without division during post-fertilization development of the fruit [18, 19].

**Figure 1.**
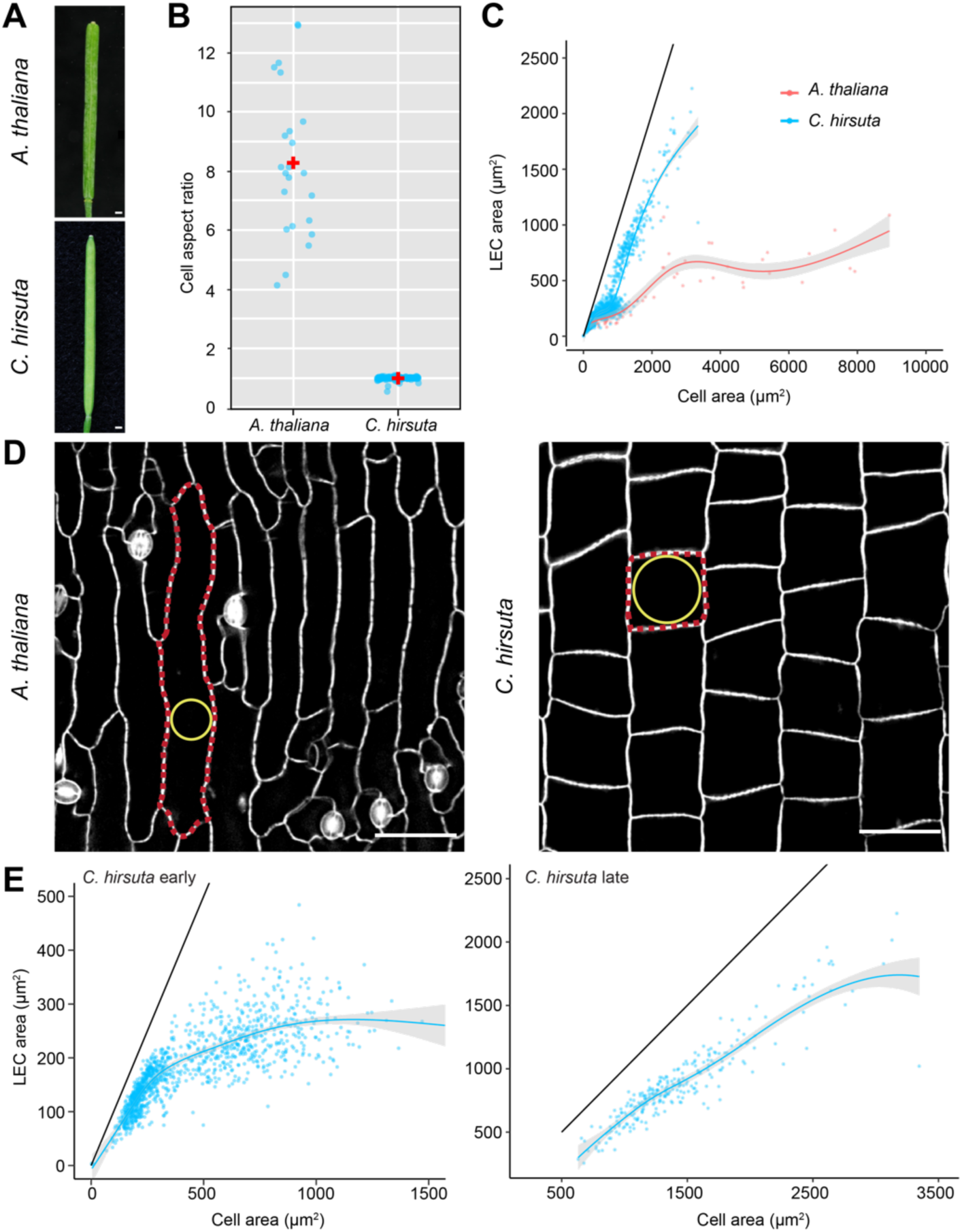
Different patterns of cell expansion underlie similar fruit shapes. **(A)** Mature fruits of *A. thaliana* and *C. hirsuta*. **(B)** Exocarp cell aspect ratio (ratio of cell length to width) in mature *A. thaliana* and *C. hirsuta* fruits; *n* = 145 cells. **(C)** Exocarp cell area (x-axis) vs the largest empty circle area (LEC, y-axis) in stage 15 – 17b fruit of *A. thaliana* (red) and *C. hirsuta* (blue). Smooth curves fitted and 95% confidence intervals (grey) shown; black line indicates perfect circle; *n* = 1568 cells. **(D)** Exocarp cells in mature fruit of *A. thaliana* and *C. hirsuta*, stained with propidium iodide; representative cell outlined (red) and largest empty circle that fits inside (LEC, yellow) indicated in each image. **(E)** *C. hirsuta* exocarp cell area (x-axis) vs LEC (y-axis) in early fruit (stage 15 – 17a, *n* = 1220 cells, upper) and late fruit (stage 17b, *n* = 258 cells, lower). Smooth curves fitted and 95% confidence intervals (grey) shown; black line indicates perfect circle. Scale bars: 1 mm (A), 50 µm (D).

Since the fruit valves grow mostly in one direction, the long thin exocarp cells found in *A. thaliana* should be sufficient to reduce the stress on the epidermal cell walls (Fig. 1D). However, enlarging the cell in two directions, as in *C. hirsuta*, creates a large open surface area (Fig. 1D), which is likely to cause the cell wall to bulge out in response to turgor and greatly increase the stress. To describe the patterns of exocarp cell expansion during different stages of fruit development in *A. thaliana* and *C. hirsuta*, we measured cell area and the area of the largest empty circle (LEC) that fits inside a cell contour (Fig. 1D). LEC has been used previously as a proxy for the magnitude of mechanical stress that accumulates in the centre of a cell surface [20]. We found that LEC area remained relatively constant as exocarp cell area increased in *A. thaliana*, (Fig. 1C), indicating that stress is likely to be minimized by this pattern of cell expansion. However, in *C. hirsuta*, LEC area increased as cell area increased (Fig. 1C), predicting a large increase in stress as cells expand. In fact, the LEC becomes twice as large in *C. hirsuta* than *A. thaliana* cells, even though cell size is much larger in *A. thaliana* (Fig. 1C-D). By analyzing *C. hirsuta* exocarp cells separately at early versus late stages of fruit development, we found that LEC area increased together with cell area only during later stages, when cells expanded in width to acquire a square shape (Fig. 1D-E). Therefore, we predict that the pattern of exocarp cell expansion during later stages of *C. hirsuta* fruit development may lead to high mechanical stress in the cell wall.

To characterize exocarp cell wall properties, and whether these change during *C. hirsuta* fruit development, we performed osmotic treatments. Turgor pressure generally increases the volume and surface area of a plant cell. Cells with isotropic material properties will expand in all directions (length, width, depth). In contrast to this, cells with anisotropic material properties, such as *C. hirsuta* exocarp cells, will expand in some directions, but contract in others [2]. We used osmotic treatments to plasmolyze exocarp cells and measure their response to increased turgor pressure at early versus late stages of fruit development (Fig. 2A-B). We quantified exocarp cell surface dimensions of plasmolyzed versus turgid cells in *35S::GFP:TUA6* plants [2], using the depolymerization of cortical microtubules as an indicator of plasmolysis [21]. During early stages of fruit development (stage 15), exocarp cells responded to increased turgor by expanding in length (+9.4% ± 0.3; mean ± S.E.M.) and contracting in width (–4.2% ± 0.4, Fig. 2A). However, during later fruit stages (stage 17b), cells contracted in length in response to increased turgor (–14% ± 0.3) and expanded in width (+14.9% ± 0.2, Fig. 2B). Thus, exocarp cells switched their response to turgor during fruit development, from length expansion to length contraction. These findings suggest that cell wall anisotropy changes during development, such that mature exocarp cells contract in length in response to turgor.

**Figure 2.**
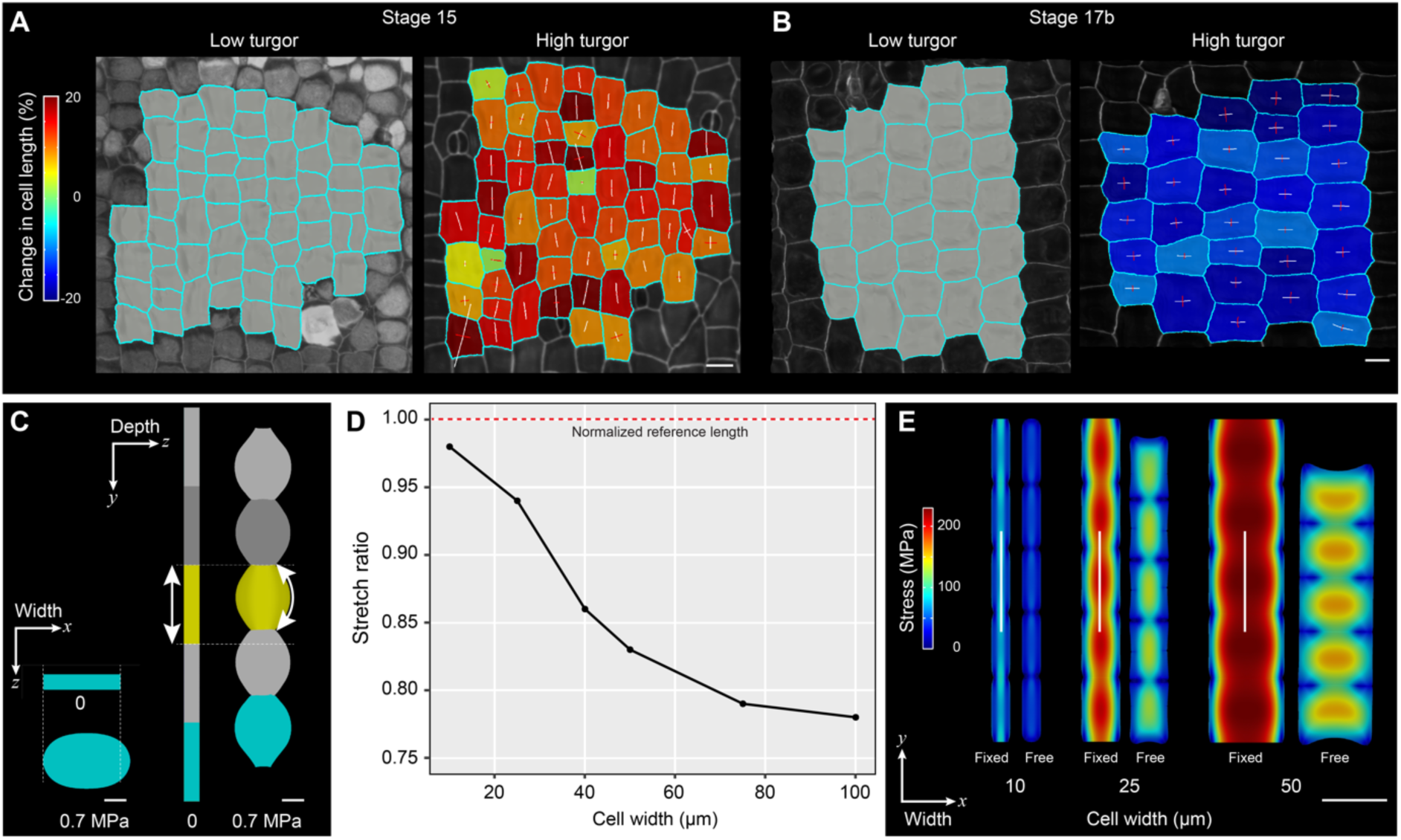
Contraction of exocarp cell length is determined by cell wall anisotropy and enhanced by cell width. **(A-B)** Osmotic treatment of stage 15 (A) and 17b (B) *C. hirsuta* fruit. Heatmaps show percentage change in exocarp cell length, relative to the long axis of the fruit, in high turgor pressure (pure water) relative to low turgor pressure (1M salt). Tensors indicate expansion (white) or shrinkage (red) of cell dimensions. Change in exocarp cell length and width shown as mean ± S.E.M. for stage 15 (*n* = 165 cells from 3 fruit) and stage 17b (*n* = 227 cells from 6 fruit). **(C-E)** FEM simulations in MorphoMechanX of cells inflated from 0 to 0.7 MPa. Material anisotropy is specified in the length direction, indicated by double headed arrow, in 50 × 50 × 10 µm (length × width × depth) cells (C). Inflated cells expand in width (cyan cell viewed end on, left) and contract in length (yellow cell viewed side on). Dashed lines indicate initial cell dimensions before inflation. High anisotropy in the length direction causes the cells to bulge in depth upon inflation (curvature of double headed arrow). Plot of stretch ratio (y-axis), measured along the length of inflated cell file when ends are free versus fixed, in six different simulations where cell width (x-axis) progressively increases (D). Length contraction compared between three simulations with different cell widths (10, 25, 50 µm, E). For each cell width, trace of Cauchy stress tensor (MPa) is compared when the inflated cell file ends are fixed (left) versus free (right). White lines indicate direction of material anisotropy in each simulation. Coordinates show the orientation of cell depth (*z*), width (*x*) and length (*y*) relative to the fruit (C, E). Scale bars: 20 µm (A-B), 10 µm (C), 50 µm (E).

We reproduced this turgor-driven contraction of mature exocarp cells using finite element method (FEM) simulations both here (Fig. 2C) and previously [2]. We found that the contraction in cell length results from the combined action of turgor pressure, which pushes the cell wall surface to expand, and high material anisotropy in the direction parallel to the cell length. This causes the cell wall to resist stretch in the length direction. When pressurized, these cells expand in width and depth, and increase their curvature much more in the length direction, causing these cell surfaces to bulge out (Fig. 2C). In this way, the cell wall expands in surface area to be in mechanical equilibrium with turgor pressure, while minimizing longitudinal stretch of the cell wall. As a consequence, the straight-line distance along the cell length contracts (Fig. 2C).

We used further FEM simulations to investigate whether expansion in cell width, which we observed during later stages of *C. hirsuta* fruit development (Fig. 1D-E), could be instrumental in building contractile force when combined with material anisotropy of the cell wall. Using inflation simulations with MorphoMechanX software (www.MorphoMechanX.org), we explored the effects of increasing cell width for a fixed combination of anisotropic material properties and pressure ([2], STARMethods). All cell files were generated with the same cell length (50 µm) and depth (10 µm), while cell width was progressively increased in each simulation (Fig. 2D). For each cell width, a file of non-pressurized cells with ends fixed in the y-direction were inflated to 0.7 MPa. When we computed the stretch ratio between the length of the cell file with free versus fixed ends (Fig. S1A), we found that increasing cell width enhanced the length contraction (Fig. 2D). The stretch ratio decreased from 0.98 to 0.78 as cell width increased from 10 µm to 100 µm, although it started to plateau beyond a cell width of 50 µm (Fig. 2D). Mechanical stress increased in the centre of the surface walls as cell width increased from 10 µm to 50 µm (Fig. 2E), particularly the longitudinal stress component that is directly connected to the pulling force (Fig. S1B). This is in agreement with the increased stress predicted by LEC area measurements of mature exocarp cells that expanded in width to acquire their final, square shape (Fig. 1D-E). A reduction in wall stress was associated with contraction of the cell file after the ends were released in the simulations (Fig. 2E). Therefore, these inflation simulations indicate that as cells become wider, they can contract more.

The relatively shallow depth of mature exocarp cells is also important for length contraction. For example, increasing cell depth from 10 µm to 50 µm elevated the stretch ratio from 0.83 to 0.92 (Fig. S1C). These cells accumulated high mechanical stress in the centre of not only surface, but also side walls (Fig. S1C). This created conflicts during the minimization of cell wall mechanical energy, which reduced the amount of length contraction in the simulation (Fig. S1C). In summary, cell shape plays an important role in amplifying exocarp cell contraction.

### Orientation of cortical microtubules and CESA3 trajectories predict growth direction

Our results from osmotic treatments and model simulations show that wider cells can contract more when inflated. To understand how exocarp cells grow into this more optimal shape, we quantified growth at cellular resolution during *C. hirsuta* fruit development. Specifically, we used time-lapse confocal microscopy to image the exocarp of fruit expressing a cortical microtubule marker (*35S::GFP:TUA6* [2]), and analyzed these confocal stacks using MorphoGraphX [22]. We began each of our time series in 7 mm long fruit, once cell division had ceased in the exocarp, and imaged three subsequent time points (48 hours (h), 96 h and 8 days (d)). We identified two phases of highly anisotropic growth during the first 48 h and the last 96 h, separated by a period of more isotropic growth from 48 – 96 h (Fig 3A). The largest amount of growth occurred during the first 48 h (Fig 3A, Fig. S3D-E). Growth anisotropy was parallel to the long axis of the fruit from 0 – 48 h and then switched to perpendicular (Fig 3A). Therefore, through this pattern of anisotropic growth, exocarp cells initially expand by elongating and then switch to growth in the width direction to acquire their final square shape.

**Figure 3.**
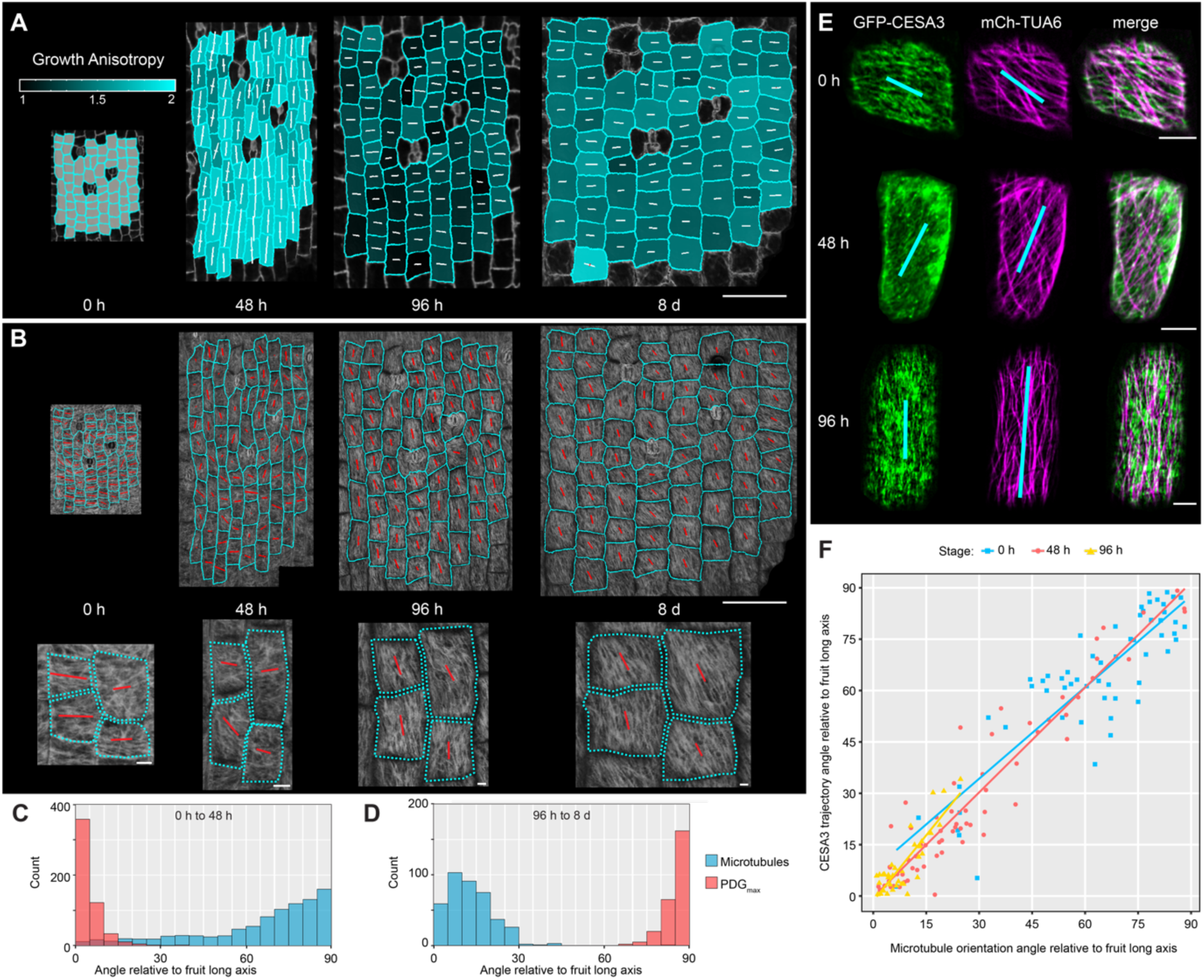
Orientation of cortical microtubules and CESA3 trajectories predict directional growth in fruit exocarp cells. **(A-B)** Time-lapse confocal imaging of *C. hirsuta* exocarp cells during 8 d of fruit development starting with 7 mm long fruit at 0 h. Growth anisotropy (ratio of PDG_max_ to PDG_min_) between consecutive time-points shown as heatmaps on second time-point (A). White tensors in each cell indicate the magnitude and direction of PDG_max_ (longer lines) and PDG_min_ (shorter lines). Cortical microtubule (CMT) marker GFP-TUA6 at 0, 48, 96 h and 8d (B). Orientation and magnitude of CMT arrays calculated with FibrilTool and shown as direction and length of red lines; cells outlined in cyan; magnified views shown below. **(C-D)** Histograms showing the principal direction of growth (PDG_max_, red) and CMT orientation (blue) as angles relative to the long axis of the fruit at 0 – 48 h (C) and 96 h – 8 d (D). PDG_max_ calculated between consecutive time-points as indicated; *n* = 535 cells (C), 253 cells (D). CMT orientation calculated at 0 h, *n* = 934 cells (C), and 96 h, *n* = 398 cells (D). **(E)** Co-localization of GFP-CESA3 and mCherry-TUA6 in exocarp cells of *C. hirsuta* fruit at 0, 48 and 96 h, starting with 7 mm long fruit at 0 h. Average projections of confocal time-lapse movies (Movies S1-S3) showing GFP-CESA3 (green), mCherry-TUA6 (magenta) and merge of both channels. Orientation and magnitude of CESA3 trajectories and CMT arrays calculated with FibrilTool and shown as direction and length of blue lines. **(F)** Scatterplot showing correlation between the angles of GFP-CESA3 trajectories and CMT orientations in fruit exocarp cells relative to the long axis of the fruit; dots indicate individual cells and lines indicate linear regressions at 0 h (blue; r = 0.87, p < 4.9 e-19, *n* = 58 cells), 48 h (red; r = 0.95, p < 4.15 e-34, *n* = 65 cells) and 96 h (yellow; r = 0.86, p < 9.87 e-11, *n* = 34 cells). Scale bars: 100 µm (A-B) 5 µm (B magnified, E).

Cells grow principally in the orthogonal direction to aligned cellulose microfibrils in the cell wall. Since the alignment of microfibrils usually mirrors the orientation of cortical microtubules (CMTs) [11], we investigated to what extent the growth anisotropy of exocarp cells was predicted by CMT orientation. The initial alignment of CMTs was perpendicular to the long axis of the fruit (0 h, Fig. 3B), giving a distribution of CMT angles skewed towards 90° (Fig. 3C). This CMT alignment was a good predictor of the principal direction of cellular growth (PDG_max_) in the orthogonal direction during the subsequent 48 h (Fig. 3C). CMT arrays began to re-align and had no principal orientation by 48 h (Fig. 3B), prior to the subsequent period of more isotropic growth (48 – 96 h Fig. 3A). By 96 h, CMT arrays were well-aligned and their orientation had switched to parallel to the long axis of the fruit (96 h, Fig. 3B), resulting in a distribution of CMT angles skewed towards 0 - 20° (Fig. 3D). This new CMT alignment accurately predicted a switch in PDG_max_ to the orthogonal direction during the subsequent 96 h (Fig. 3D). This CMT alignment parallel to the fruit long axis was maintained through to the last time point at 8 d (Fig. 3B). Therefore, CMT orientation is highly predictive of growth anisotropy in the fruit exocarp.

However, CMT orientation is only a proxy for cellulose microfibril alignment. Therefore, we investigated whether cellulose synthase (CESA) proteins tracked along CMTs during exocarp cell growth. To address this question, we co-localized CESA3 (*pCESA3::GFP:CESA3*) and CMTs (*pUbi10::mCherry:TUA6*) in exocarp cells of *C. hirsuta* fruit during 96 h of development (Fig. 3E). We visualized CESA3 particles tracking along CMTs (Supplementary movies S1-S3) and measured the orientation of CMTs and CESA3 trajectories per cell in average projections of these time-lapse movies (Fig. 3E). The orientation of CMTs and CESA3 trajectories were highly correlated at each time point (Fig. 3F), and showed the same switch from transverse to longitudinal orientations that we observed previously (Fig. 3B-E). Therefore, CMT alignment dictates the orientation in which new cellulose microfibrils are synthesized in exocarp cells of *C. hirsuta* fruit. Consequently, CMT realignment in mature exocarp cells produces a change in both cell wall anisotropy and growth direction.

### Dynamics of growth and tension

By switching the direction of cell wall reinforcement and growth, exocarp cells grow to a more optimal shape to build contractile tension. But, how is it possible that growth, a process that relaxes stress in the cell wall, generates more contractile tension in the valve?

To explore this apparent paradox, we performed growth simulations using MorphoMechanX software (STARMethods, Supplementary Information). We used a turgor-driven, strain-based growth model, where growth depends on the amount of elastic strain on the cell wall, resulting from turgor pressure, and irreversible cell wall extensibility [9, 17, 23]. When we added this growth model to cell file simulations with material anisotropy in the length direction, we found that maximal strain, and hence growth, was oriented in the width direction (Fig. 4A). This matches the growth we observed in cell width when the deposition of cellulose microfibrils was reoriented to the length direction in exocarp cells (Fig. 3).

**Figure 4.**
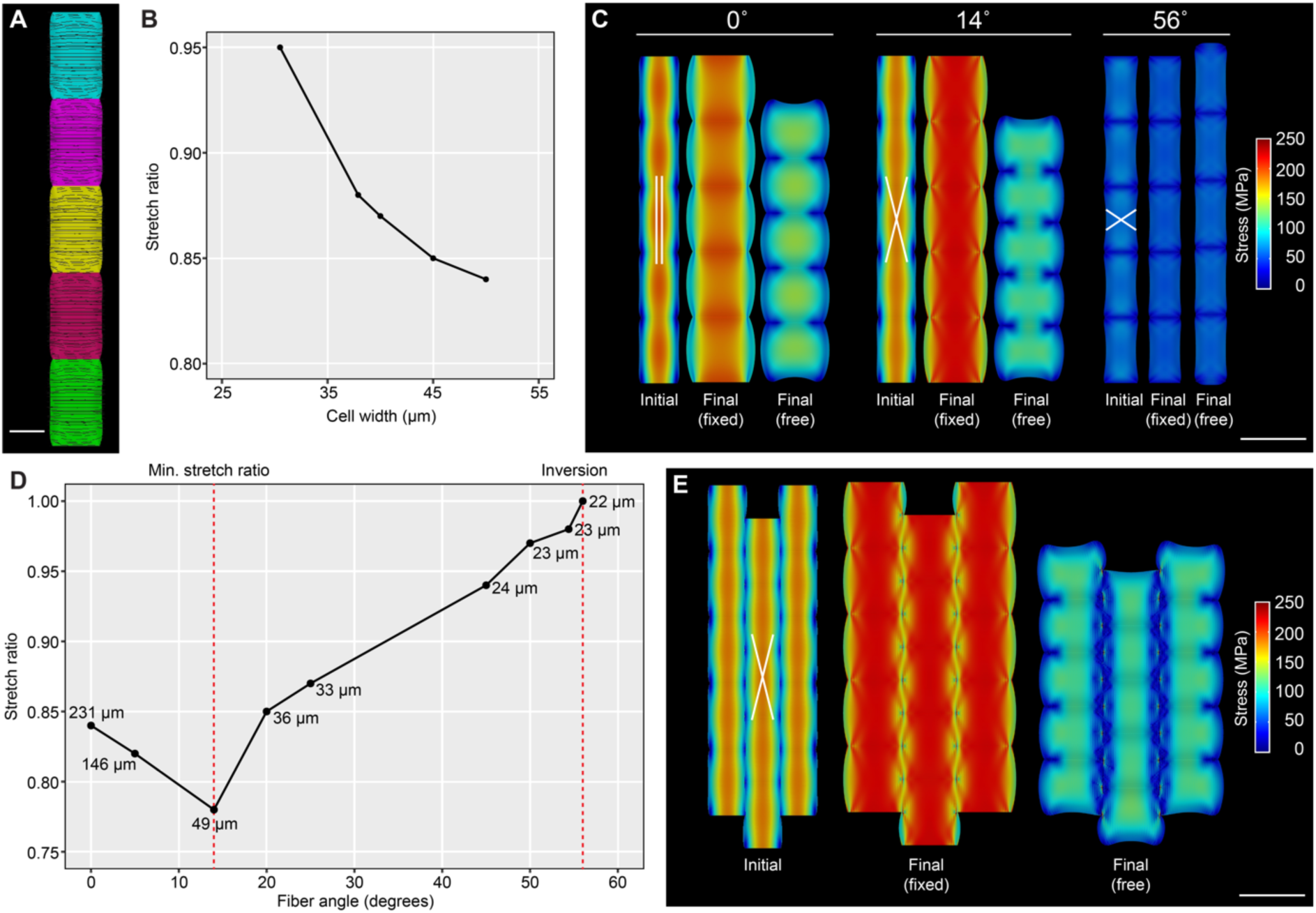
Interaction of growth and cell wall anisotropy determines cell length contraction. All FEM growth simulations performed in MorphoMechanX using cell templates of dimensions 50 x 26 x 10 µm (length, width and depth) inflated to 0.7 MPa. **(A-B)** Strain-based growth model comprising transversely isotropic material reinforced with longitudinal fibers of 4000 MPa stiffness. Direction of maximal strain, and therefore main growth direction, indicated by black lines in a file of inflated cells before growth (A). Plot of stretch ratio (y-axis), measured along the cell file length when ends are free versus fixed, at intermediate steps of the growth simulation as cell width (x-axis) increases to a final target of 50 µm (B). **(C-E)** Multi-layer growth model comprising an isotropic layer undergoing strain-based growth joined with two layers of stiff fibers (2000 MPa each layer) that do not respond to growth. Schematic of fiber orientation in each of the two layers is shown on initial cell file. Shown for each simulation: a file of inflated cells with fixed ends before strain-based growth starts (Initial), at the end of growth (Final, fixed) and once the ends of the cell file are released (Final, free). Length contraction and stress compared between three simulations with either 0°, 14° or 56° fiber angles, with respect to vertical (C). Stress ratios: 0.84 (0°), 0.78 (14°), 1.005 (56°). Phase-space plot of stretch ratio (y-axis), measured along the cell file length when ends are free versus fixed, in simulations with different fiber angles (x-axis) from 0° through to 56° (D). Stretch ratios computed at 50 µm cell width or maximal width if less than 50 µm. Maximal cell width achieved through growth is annotated on the plot for each simulation. Fiber angles of 14° (minimal stretch ratio) and 56° (inversion) are highlighted by red dashed lines. Growth simulation in a block of cells with 14° fiber angle (E). Stress increases during growth to a maximum cell width of 41.4 µm (compare Initial versus Final, fixed) and drops once the cell file ends are released (stretch ratio 0.82). Heatmaps display trace of Cauchy stress tensor. Scale bars: 20 µm (A), 50 µm (C, E).

To investigate how this growth could lead to an increase in contractile tension, we designed a series of simulations with a file of cells geometrically comparable to exocarp cells starting growth in the width direction. Cell size was set to 50 × 26 × 10 µm (length, width and depth) and other mechanical parameters were assigned identical to previous inflation simulations (STARMethods). Turgor pressure was assumed to remain constant [2] and strain-based growth was set uniformly. The simulation cycle consisted of (i) cell pressurization at a fixed value that is maintained throughout the simulation, (ii) a growth step, in which accumulated mechanical strain is used to proportionally grow the cell wall, and (iii) recalculation and update of mechanical equilibrium (STARMethods). Steps (ii) and (iii) were reiterated until cells reached a target width of 50 µm, which matches our measurements of mature exocarp cell width (Fig. 1D, Fig. 3A). We fixed the ends of the cell file to mimic the constraint exerted on the growing exocarp by its physical attachment to the rest of the fruit.

Following our hypothesis that the change in cell shape due to growth causes an increase in contractile tension, we assessed the effect of growth on cell length contraction. To do this, we saved the grown cell file template at intermediate growth steps with all the relevant information about its mechanical state, including the mechanical and growth properties assigned, its deformation status and reference configuration. At the end of the full growth simulation, the saved intermediate meshes during the growth process were retrieved, the cell file ends released, and the mechanical equilibrium was recomputed. In this way, we could measure the stretch ratios at intermediate growth steps in the process (Fig. 4B).

We tested two growth scenarios where we allowed almost none (Fig. 4B), or some (Fig. S2A), relaxation of cell wall stress in the stiffer, length direction. This was done by modulating the yield threshold of the cell wall, which has to be exceeded to activate growth; a higher threshold of 0.1 in the first scenario and a lower threshold of 0.01 in the latter (STARMethods and Supplementary Information). As the soft direction of the wall is stretched beyond both of these thresholds, the yield criterion will only significantly affect growth in the stiff direction. As cell width increased through growth from approximately 30 µm to 50 µm, we found that the stretch ratio of the cell file decreased from 0.93 to 0.89 when the yield threshold was low (Fig. S2A, Table 1). At the higher yield threshold, the stretch ratio decreased even further to 0.84 (Fig. 4B, Fig. S2A, Table 1). These results demonstrate that although growth is a stress-release process, the release of stress in the width direction can increase stress in the length direction, by changing cell geometry and increasing the stress associated with a larger cell surface area. This requires that the relaxation of wall stress in the length direction is inhibited, in this case by significant reinforcement of cellulose microfibrils in that direction, which reduces the strain below the yield threshold.

**Table 1.** Summary of stretch ratios from FEM growth simulations, related to Fig. 4 and Fig. S2. Stretch ratios measured along the cell file length for different growth model simulations after inflation (before growth) and after growth (when ends are free versus fixed). Stretch ratios were computed after growth achieved a target cell width of 50 µm or else the maximal cell width achieved if less than 50 µm.

| Growth model | Stretch ratio after<br>inflation | Stretch ratio after<br>growth |
| --- | --- | --- |
| Transversely isotropic material, low strain threshold (0.01, Fig. S2A) | 0.93 | 0.89 |
| Transversely isotropic material, high strain threshold (0.1, Fig. 4B, Fig. S2A) | 0.93 | 0.84 |
| Multi-layer, cell file, 0° fiber angle (Fig. 4C) | 0.94 | 0.84 |
| Multi-layer, cell file, 14° fiber angle (Fig. 4C) | 0.94 | 0.78 |
| Multi-layer, cell file, 56° fiber angle (Fig. 4C) | 1 | 1.02 |
| Multi-layer, cell block, 0° fiber angle (Fig. S2F) | 0.93 | 0.86 |
| Multi-layer, cell block, 14° fiber angle (Fig. 4E) | 0.93 | 0.82 |

In summary, the stretch ratio of 0.84 obtained for a simulated file of growing cells (Fig. 4B, Table 1), is comparable to the empirical value of 0.8 reported previously [2]. Therefore, material anisotropy, specified as longitudinally aligned fiber reinforcement, combined with turgor-driven growth and limited fiber relaxation, can explain the generation of contractility as a file of cells grows in width.

### A cross-lamellate pattern of cellulose microfibrils enhances contractility

Like many biological processes, the longitudinal alignment of CMTs and associated CESA3 trajectories observed in mature exocarp cells are distributed around a mean, both within and between cells (Fig 3B-E). CMT arrays are dynamic and unstable, reorganizing on a scale of minutes [24, 25], and these arrays guide only the most recently added layer of cellulose microfibrils in the cell wall. This leads to heterogeneity in microfibril orientation between different cell wall layers, contributing to a cross-lamellate pattern [26–28]. This type of pattern is observed in mature exocarp cells of *C. hirsuta* fruit where longitudinally aligned cellulose microfibrils intersect at acute angles [2]. For these reasons, we performed a new series of simulations to explore whether deviations from a parallel fiber orientation can affect the process of building contractility in growing exocarp cells.

To vary the orientation of fiber reinforcement in the cell wall, we designed a multi-layer model. The cell wall is treated as a continuum in all of our models, rather than an explicit description of discrete fibers. Therefore, we used multiple layers to ascribe distinct orientations of fiber reinforcement within a growing, composite material. This multi-layer model comprised an isotropic layer that grows, and two different fiber-dominated layers, representing the averaged contribution of cellulose microfibrils oriented at distinct angles in each layer, which do not grow (Fig. S2B). The anisotropic component of the previous growth model was essentially split between the two fiber-dominated layers (parameter details in STARMethods). These fiber-dominated layers are prevented from growing due to the high density of oriented cellulose microfibrils, which, according to our previous results (Fig. 4B), will not be able to stretch beyond the yield threshold that initiates growth. The isotropic layer represents the cell wall matrix with some amount of randomly oriented cellulose microfibrils, which undergoes turgor-driven, strain-based growth [9, 17, 23]. The three layers are geometrically joined, meaning that their mesh nodes are shared, their displacement is unique, and the nodal forces moving the points towards their equilibrium configuration are computed as the sum of the forces provided by each layer (Supplementary Information, STARMethods). In this way, the three layers can be treated as a composite material (Fig. S2B).

In order to compare this multi-layer model with our previous growth model, we assigned a common fiber direction parallel to the longitudinal cell file in both fiber layers. Under a hypothesis of full compressibility (Poisson ratios equal to zero), where the two modeling approaches are theoretically comparable (STARMethods), the multi-layer model produced the same stress pattern and deformation as the previous growth model (Fig. S2E). We also tested the effect of imposing symmetry on the fiber angles in each layer. Comparable results were produced with asymmetric fiber angles, although the cell file twisted when its ends were released (Fig. S2C). Therefore, symmetric angles allowed a simpler computation of stretch ratios.

We performed simulations with two symmetric families of fibers at increasing angles relative to the vertical direction (Fig. 4C-D). We computed the stretch ratio at the end of the growth process, when cells reached a target width of either 50 µm or the maximal width attained if less than 50 µm. As before, we calculated the stretch ratio by comparing the final cell file length with free versus fixed ends (Fig. 4C, Fig. S1A). To test the effect of fiber angle on stretch ratio, we varied the angle from 0° up to 56° with respect to vertical (Fig. 4D). We found that the stretch ratio reached a minimum around a fiber angle of 14° and increased at angles above 14° until an inversion point of 56° (Fig. 4D). Interestingly, this point of inversion at 56° is in agreement with analytical predictions for a cylinder under pressure that is reinforced by two populations of fibers at symmetric angles to vertical [29].

In simulations where the fiber angles were 0° or 14°, we found a large reduction in stress was associated with contraction of the final cell file once the ends were released (Fig. 4C). The stretch ratio was 0.84 when the fiber angle was 0° (Fig. 4C, Table 1). This value is identical to the previous growth model with parallel fiber reinforcement, grown to 50 µm cell width (Fig. 4B, Table 1), providing further evidence that the two modeling approaches are comparable. However, as the fiber angle changed from parallel to 14°, with respect to vertical, we found that the stretch ratio of the cell file decreased from 0.84 to 0.78 (Fig. 4C, Table 1). Since both cell files had the same stretch ratio before growth (0.94, Table 1), this suggests that the higher contractility of the cells with crossed fibers (14°) is actively reached through growth. At a fiber angle of 56°, growth was negligible and we found an inversion in the stretch behavior as the cell file expanded in length with a stretch ratio reaching 1.02 (Fig. 4C, Table 1).

The reason that fibers in a crossed pattern (14°) can achieve a lower stretch ratio lies in the additional tension exerted on the fibers during growth. The expansion in cell width due to growth of the isotropic layer, exerts an additional elastic stretch on the fiber layers when the fibers are tilted rather than orthogonal to the direction of growth. This stretch in the direction of growth is transmitted along the tilted fiber direction and partially translated into additional pulling force (Fig. S2D). Due to this interaction with growth, our findings suggest that, rather than being noise, crossed patterns of cellulose microfibrils can actually enhance the process of building contractile tension in growing exocarp cells.

Our findings also revealed the influence of increasing fiber angles on the maximum cell width achieved by growth. At fiber angles closest to parallel (0°), cells achieved the highest growth in width, and at angles from 14° and above, cells could no longer reach a width of 50 µm (Fig. 4D). At these angles, the fibers restrict the amount of transverse strain generated for growth in width. Therefore, in our simulations, when fibers were crossed and offset at a slight angle to vertical, this limited the final cell width attained by growth, providing an intrinsic mechanism to halt the growth process.

To verify that our results are not limited to the simplified case of a single cell file, we implemented our multi-layer growth model in a block of cells which more realistically represents the growth of cells in the context of exocarp tissue (Fig.4 E, Fig. S2F, STARMethods). A large reduction in stress was associated with contraction of the cell block length once the ends were released after the growth process (Fig. 4E, Fig. S2F). The stretch ratio of the central cell file in this block was 0.82 at a fiber angle of 14°, slightly higher than the ratio of 0.78 achieved in a single cell file (Table 1). The block configuration constrained the growth in width of these central cells such that they reached only 41.4 µm (Fig. 4E). Even with this constraint, the stretch ratio of the central cell file with fibers offset at 14° (0.82, Fig. 4E) was lower than a cell file with parallel fibers in both a block configuration (0.86, Fig. S2F) or as a single file (0.84, Fig. 4C) (Table 1). Thus, our findings can be generally applied to a tissue context.

The stretch ratio is a simple and precise measure of the amount of contraction occurring in the exocarp both *in planta* and *in silico*. However, we were also interested in the force produced by exocarp contraction since this is fundamental to the explosive nature of seed dispersal in *C. hirsuta*. We previously used empirical extensometer experiments to measure the force required to extend coiled fruit valves, and then compute the force exerted by the exocarp layer before explosion [2]. To compare our simulations with these results, we measured the force exerted at the ends of a cell file and extrapolated this to the pulling force for the whole exocarp layer (valve width 1 mm, Table 2). Our results fell within the range of previously determined forces (Table 2). We found that simple inflation simulations using tissue templates made of wider cells (~50 versus ~29 µm final cell width, Table 2) contracted more and produced more exocarp layer pulling force (Table 2). We found a similar increase in cell contraction and pulling force when cell width increased to ~50 µm, from an initial width of 26 µm, in growth simulations (Table 2). Interestingly, we found that growing cells with a crossed pattern of fibers, offset at 14°, produced the greatest pulling force (Table 2). This result indicates that if cells reach an optimal width through growth, they contract more and pull with greater force. This effect depends on the interaction of growth with crossed fibers (Table 2, Fig. S2D). In summary, our simulations show that growth actively generates more contraction at the cell level and more pulling force at the tissue level, which contributes to the mean velocity of the coiling valve.

**Table 2.** Simulated exocarp pulling force. Force exerted at the ends of an exocarp cell layer after growth or inflation simulations. Final cell width, stretch ratio, and corresponding pulling force extrapolated from a cell file to a cell layer (valve width: 1 mm), are measured for the different multi-layer model templates indicated. Fiber angle in the multi-layer model is either crossed (14° angle) or parallel (0° angle) as indicated. Exocarp pulling force previously computed from extensometer experiments for comparison: average = 37 mN, max = 75 mN [2].

| FEM simulation, cell template:<br>width × length × depth (μm) | Final cell width<br>(μm) | Stretch ratio | Exocarp pulling<br>force (mN) |
| --- | --- | --- | --- |
| Crossed fibers (14° angle) |  |  |  |
| Inflation, 26 × 50 × 10 | 29 | 0.94 | 35.6 |
| Inflation, 45 × 50 × 10 | 50 | 0.84 | 44.0 |
| Growth, 26 × 50 × 10 | 49 | 0.78 | 62.9 |
| Parallel fibers (0° angle) |  |  |  |
| Inflation, 26 × 50 × 10 | 29.4 | 0.94 | 32.4 |
| Inflation, 45 × 50 × 10 | 50.6 | 0.85 | 44.1 |
| Growth, 26 × 50 × 10 | 51.6 | 0.84 | 39.9 |

### Cortical microtubules (CMTs) contribute to explosive valve coiling

Our findings predict that explosive valve coiling depends on exocarp CMTs, since these control the specific orientation of cellulose microfibrils in the cell wall, and the consequent cellular growth pattern. To test this prediction, we performed microtubule perturbation experiments. Treating *C. hirsuta* fruit with the microtubule depolymerizing drug oryzalin, resulted in non-explosive fruit where the valves failed to coil (Fig. S3A). However, it was not possible to ascribe this effect exclusively to exocarp cells, since microtubules were depolymerized in all cell types of the fruit in these experiments.

To achieve more precise spatial and temporal resolution, we used a genetic system to depolymerize microtubules via inducible expression of a truncated version of the atypical tubulin kinase PROPYZAMIDE-HYPERSENSITIVE 1 (PHS1ΔP) [30]. We expressed PHS1ΔP in exocarp cells using an epidermal-specific promoter (*pML1::GR-LhG4/pOp6::PHS1ΔP:mCherry*) and verified that cortical microtubules were depolymerized specifically in response to dexamethasone and only in epidermal layers of the fruit (Fig. S3B-C). For example, the mechanically important endocarp *b* cell layer of the fruit valve was unaffected in these transgenic lines (Fig. S3C). Using live confocal imaging of *pML1::LhGR>>PHS1ΔP:mCherry; 35S::GFP:TUA6* plants, we imaged exocarp cells in 7 mm long fruit, immediately before dexamethasone treatment and at three subsequent time points (48 h, 96 h and 8 d, Fig. 5). The initial alignment of exocarp CMTs was perpendicular to the long axis of the fruit (pre-induction, Fig. 5A). Fruits were treated with dexamethasone at 0, 24 h, 48 h, 96 h and then every 48 h until day 11. Imaging at 48 h, 96 h and 8 d, showed that microtubule depolymerization was maintained throughout the time-lapse experiment (Fig. 5A).

**Figure 5.**
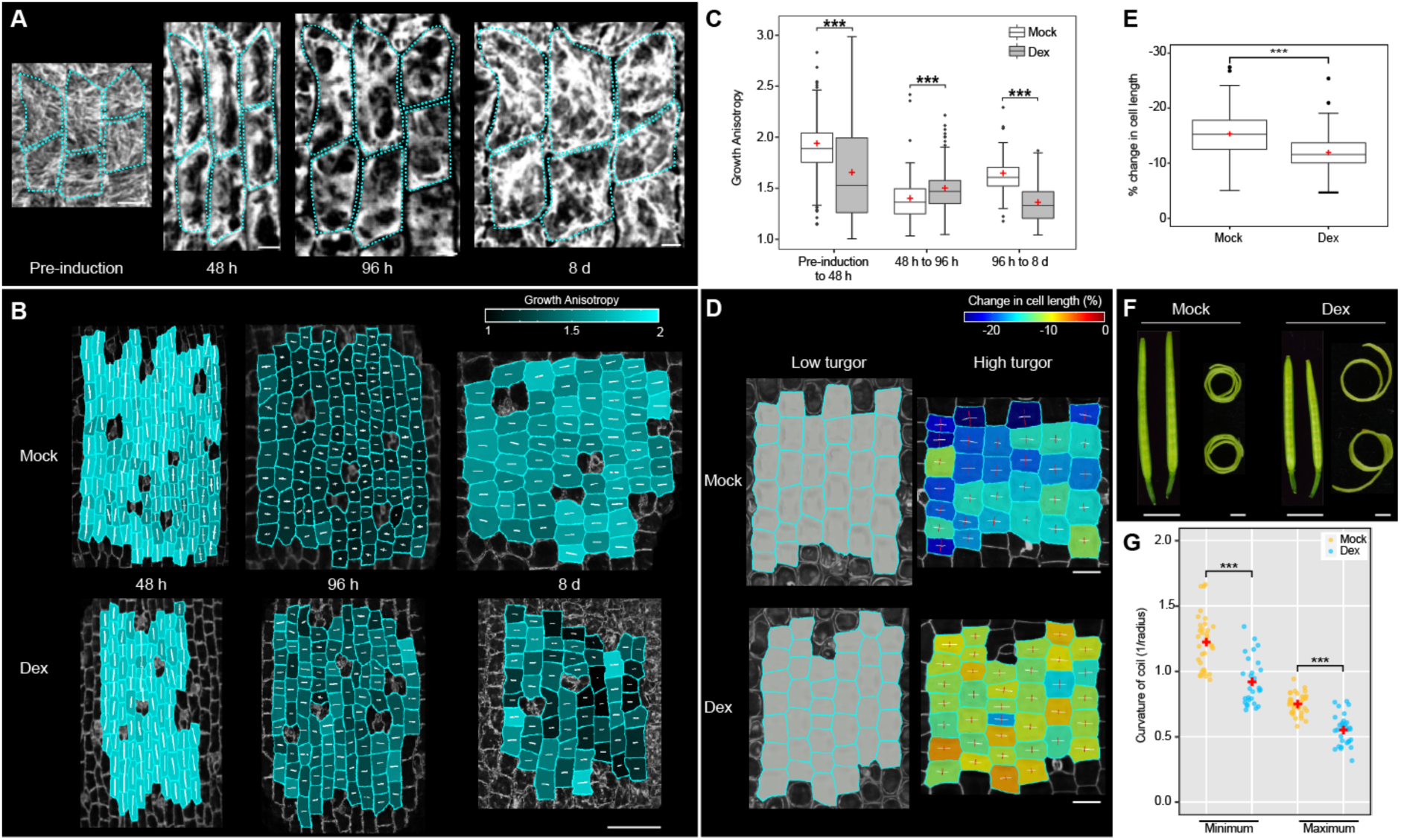
Cortical microtubules in fruit exocarp cells contribute to explosive valve coiling. Time-lapse confocal imaging of *C. hirsuta pML1::LhGR>>PHS1ΔP:mCherry; 35S::GFP:TUA6* exocarp cells during 8 d of fruit development starting with 7 mm long fruit at 0 h. **(A)** Cortical microtubule (CMT) marker GFP-TUA6 immediately before (pre-induction) and 48, 96 h and 8d after dexamethasone treatment. Individual cells outlined in cyan. **(B-C)** Growth anisotropy (ratio of PDG_max_ to PDG_min_) between consecutive time-points in mock- and dexamethasone-treated (Dex) fruit. Shown as heatmaps on second time-point; white tensors in each cell indicate the magnitude and direction of PDG_max_ (longer lines) and PDG_min_ (shorter lines) (B). Shown as boxplots (C), *n* ≥ 260 cells from 4 fruit per time-point (Mock), 262 cells from 5 fruit per time-point (Dex). **(D-E)** Osmotic treatment of mock- and dexamethasone-treated fruit. Percentage change in exocarp cell length, relative to the long axis of the fruit, in high turgor pressure (pure water) relative to low turgor pressure (1M salt); shown as heatmaps, tensors indicate expansion (white) or shrinkage (red) of cell dimensions (D), and boxplots (E), *n* = 195 cells from 4 fruit (Mock), 164 cells from 3 fruit (Dex). **(F-G)** Mock- and dexamethasone-treated fruit and coiled valves, 11 d after treatment (F), and scatterplot showing minimum and maximum curvature of coiled valves (1/radius) (G), *n* = 35 valves (Mock), 31 valves (Dex). All plots indicate mean (red cross) and significant differences (***) at *p* < 0.001 using a Wilcoxon signed-rank test or Student’s *t*-test. Scale bars: 10 µm (A), 100 µm (B), 50 µm (D), 1 mm (F).

Following microtubule depolymerization, exocarp cell growth was slower and more isotropic (Fig. 5B, Fig. S3D-E). In particular, dexamethasone treatment significantly reduced the anisotropic growth in width observed in mock-treated fruit (96 h – 8 d, Fig. 5B-C, Fig. S3F-G). To test whether cell wall anisotropy was disrupted by microtubule depolymerization, we used osmotic treatments. Mature exocarp cells of dexamethasone-treated compared to mock-treated fruit, contracted significantly less in length in response to increased turgor (Fig. 5D-E). Thus, in the absence of CMTs to guide the orientation of newly deposited cellulose microfibrils, cell wall anisotropy was reduced.

To analyze the effect of microtubule depolymerization on explosive valve coiling, we triggered fruit to explode 11 days after the first dexamethasone treatment. The detached valves from dexamethasone-treated fruit coiled more loosely, with reduced curvature, compared to the tightly coiled valves from mock-treated fruit (Fig. 5F-G). Thus, microtubule dynamics contribute to the patterns of cellular growth and cell wall anisotropy that determine exocarp cell contraction and underpin explosive valve coiling.

## Discussion

Exploding seed pods in *C. hirsuta* harness general properties of cellular growth in living tissues to actively contract. We identified a key role for cortical microtubule dynamics in this process. A switch in microtubule alignment reorients the direction of cellulose microfibril synthesis, changing growth direction and consequently cell shape. The large, square surface walls of these cells, reinforced by aligned cellulose microfibrils, bulge out in response to turgor pressure and pull in the length direction. This puts the outer exocarp layer of the fruit valve under tension and the lignified inner layer under compression, resulting in the build-up and storage of potential elastic energy that powers explosive seed dispersal. Thus, by combining computational modeling with biological experiments, we could explain how specific cellular features cause the tissue level mechanics underpinning explosive dispersal.

The advance we make here is to explain how exocarp cells acquire the ability to contract through growth. We previously explained exocarp cell contraction in terms of its elastic behaviour [2], but not the dynamics of growth and tension. Since elongation of the fruit occurs before contractile tension develops in the valve [2], the contribution of growth to this process was unclear. Specifically, it was unclear how growth, which is a stress relaxation process, could generate tension.

The key is in the cell geometry. Exocarp cells initially grow by elongating, but later in fruit development they switch to grow in width. As they widen, the surface and bottom walls of these relatively shallow cells increase their surface area, causing mechanical stress to accumulate. We could measure this stress directly in inflation simulations (Fig. 2E) and by proxy using the size of the largest empty circle (LEC, Fig. 1C-E). The accummulated stress in these anisotropic walls causes them to bulge out, shortening the cell. Therefore, the role of growth is to allow cells to transition to a more optimal shape to contract and generate pulling force. However, this alone would not be sufficient to produce a significant amount of contraction since growth occurs by stress relaxation. It is important that the cell wall resists yielding to stretch in the direction of cellulose microfibril reinforcement in order not to lose its longitudinal elastic tension under the action of growth.

A second mechanism, that acts in concert with cell shape, is the cross-lamellate pattern of cellulose microfibrils in the cell wall. Growth in the width direction exerts an extra stretch on the fibers, making them pull harder in the length direction (Fig. S2D). In this way, growth becomes an active element that increases contractility beyond just the effect on cell geometry. As such, the exocarp behaves as a novel analogue of a McKibben actuator, increasing its ability to contract and exert pulling force through growth. This observation has potential applications in the field of biomimetics in connection with soft-actuator design.

A key developmental event in this contraction process is the switch in microtubule orientation. Therefore, it is of interest to understand how microtubules reorient in exocarp cells of *C. hirsuta* fruit. Much attention has focussed on the alignment of microtubules with the direction of maximal tension in plant tissues [31–35]. For this reason, microtubules are proposed to spontaneously orient with the direction of maximal tensile stress [36]. However, in fruit exocarp cells, microtubules reorient to align with the direction of minimal rather than maximal tensile stress. Based on both the geometry and cell wall anisotropy of exocarp cells prior to microtubule reoriention, maximal tensile stress is predicted to be oriented in the transverse direction [9, 23]. Yet microtubules reorient to the longitudinal direction, suggesting that a developmental signal possibly overrides other stress sensing mechanisms to orient microtubules in these cells. Identifying such a signal and understanding how exocarp microtubules switch orientation will be an important follow-up to this study.

By using a genetic system to perturb microtubules, which is tissue-specific and inducible [30], we showed that cortical microtubules in the exocarp cell layer contribute to cell wall and growth anisotropy, and ultimately to explosive coiling of the fruit valves. However, this perturbation did not abolish explosive valve coiling to the same extent as oryzalin treatment. Since oryzalin treatment is not tissue-specific, it also disrupted secondary cell wall patterning in the endocarp *b* cell layer of the valves, which is known to be critical for explosive coiling [2]. Therefore, the effect of exocarp microtubules on valve coiling was confounded in these drug treatments. However, the results from our genetic approach may still underestimate the role of exocarp microtubules. Since cellular growth of the exocarp cell layer is constrained by its physical attachment to underlying tissues, it is not free to grow fully isotropically in response to microtubule depolymerization. Understanding how microtuble reorientation is genetically regulated during exocarp cell growth will provide new approaches to understand the contributuion of exocarp microtubules to explosive valve coiling.

We noticed a complete loss of CESA3 particles from the plasma membrane of exocarp cells soon after the 96 h timepoint in our experiments (Fig. 3E). A similar depletion of primary cell wall CESA proteins during xylem vessel development was shown to be caused by internalization and reduced delivery of CESAs to the plasma membrane [37]. This suggests that subsequent expansion of exocarp cells may continue without active cellulose biosynthesis or possibly with an altered composition of CESA proteins. These findings also highlight the importance of co-localizing cortical microtubules and CESA proteins in different cell types and at different stages of development, since microtubule alignment cannot always be assumed to indicate active cellulose microfibril synthesis.

Our simulations indicate that a cross-lamellate pattern of cellulose microfibrils can enhance the active contraction of growing exocarp cells. Although it might have been possible to address this question using a classical modelling approach with two families of fibers [38], the multi-layer formulation that we designed here has several advantages including: (1) the intuitive meaning of mechanical parameters assigned to each layer; (2) the ability to prescribe individual growth rules to each layer; (3) computational ease. For example, it is possible to extend the model to any number of layers without additional complexity or significant computational cost. For these reasons, we suggest that this type of multi-layer modeling approach has potential applications beyond the scope of this study. For example, it opens up exciting possibilities to model the mechanics of plant cell walls [39]. By discriminating newly deposited layers from progressively aging layers, it allows testable hypotheses to be formulated about the differential contribution of these layers to cell mechanics, growth and morphogenesis.

In summary, exocarp cells harness mechanical forces associated with turgor-driven growth to contract and generate tension in exploding seed pods of *C. hirsuta*. This illustrates how a novel mechanical trait can emerge from tinkering with existing components during evolution [40].

## Supporting information

STARMethods

## Acknowledgements

We thank D. Moulton and M. Tsiantis for comments, M. Majda for preliminary observations, A. Sampathkumar and A. Maizel for sharing plasmids, and A. Emonet for graphic designs. This work was supported by Deutsche Forschungsgemeinschaft FOR2581 Plant Morphodynamics grants to A.H. and R.S.S., and support from University of Zurich Forschungskredit to G.M.

