## Supplementary Figures S1-S3 for "Growth and tension in explosive fruit"

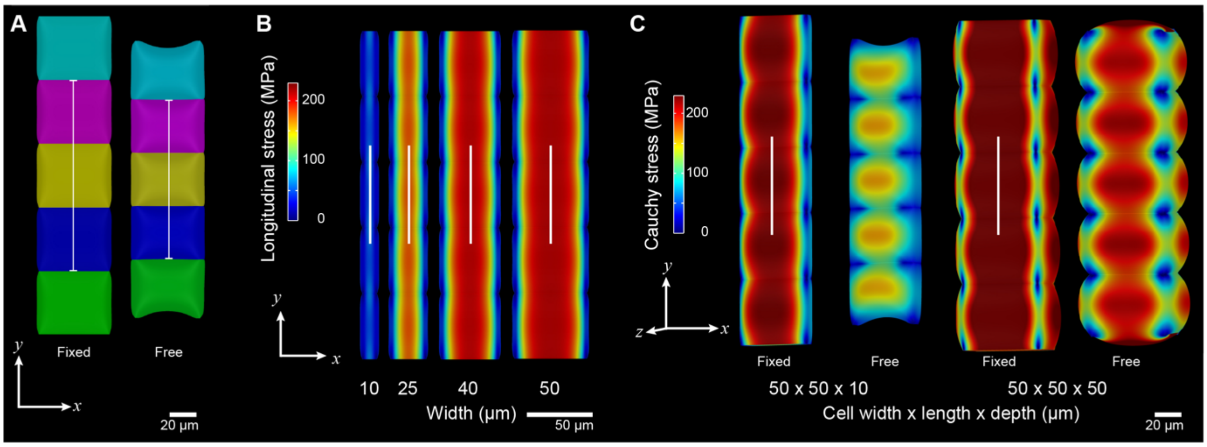


**Figure S1. FEM inflation simulations, related to Fig. 2. (A)** Stretch ratio measured along the length of an inflated cell file when ends are fixed (left) versus free (right); the two outermost cells in the file are excluded to minimize the effect of boundary deformation. **(B)** Longitudinal stress (MPa), which is parallel to the direction of contraction and connected with the average pulling stress (pulling force over cell wall cross sectional surface), compared between four inflated cell files with different cell widths (10, 25, 40, 50 μm). All files have 50 × 10 μm (cell length, depth). **(C)** Length contraction (after fixed ends of an inflated cell file are freed) compared between two cell files that differ in depth: left, 50 × 50 × 10 μm and right, 50 × 50 × 50 μm (cell width, length, depth), shown in tilted view. Heatmap displays trace of Cauchy stress (MPa). Length contraction is greater for the file of more shallow cells (left), even though the accumulated stress during inflation is higher for the file of deeper cells (right). White lines indicate direction of material anisotropy in each simulation (B-C).

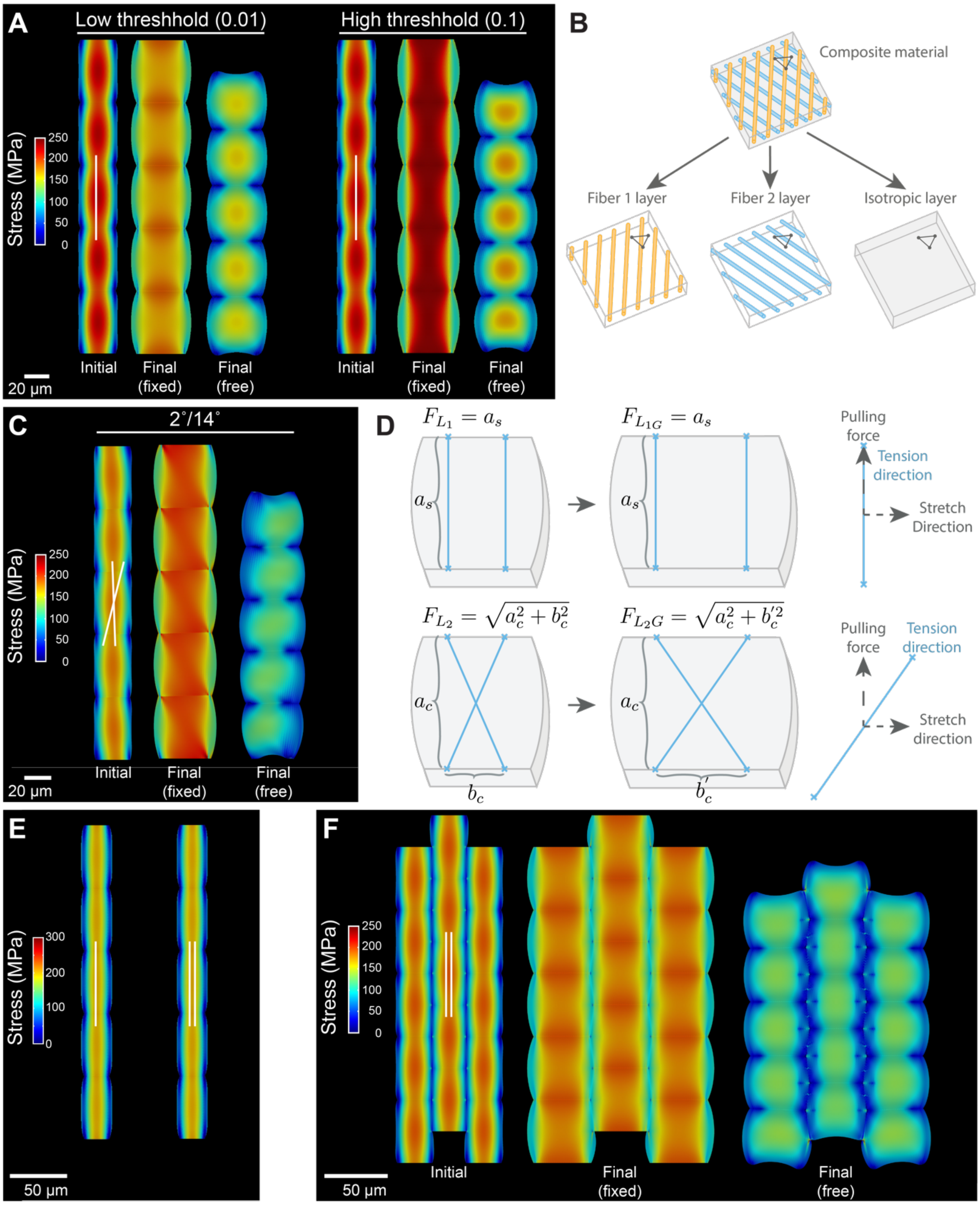


**Figure S2. FEM growth simulations, related to Fig. 4. (A)** Length contraction compared between two growing cell files with different yield thresholds for strain-based growth (0.01 versus 0.1). Shown for each simulation: a file of inflated cells with fixed ends before strain-based growth starts (Initial), at the end of growth (Final, fixed) and once the ends of the cell file are released (Final, free). A higher threshold for strain-based growth results in greater contraction in length: stretch ratio 0.84 (right) versus 0.89 (left). **(B)** Schematic of multi-layer model comprising two fiber-dominated layers (with different fiber orientations indicated in blue and orange) and one isotropic layer. Material and growth properties can be independently assigned for each layer, but the layers are geometrically fused (indicated by shared triangle). **(C)** Growth simulation of multi-layer model, identical to Fig. 4C-D, but with two non-symmetric fiber orientations (2º and 14º with respect to vertical). Initial and final stages of the growth simulation, with cell file ends fixed versus free (stretch ratio 0.825), are shown as above. This indicates that cell length contraction is enhanced by crossed fibers, even without the symmetry constraint, but contraction is maximized by having both families of fibers at the optimal angle of 14º (compare Fig. 4C-D). Note how the stress pattern is clearly non-symmetric with respect to vertical after growth (Final, fixed), and how the cell file twists once its ends are released (Final, free). **(D)** Schematic showing the principle by which a crossed pattern of fibers in the cell wall (lower panel) can cause a higher accumulation of contractile tension during growth, as compared to fibers aligned parallel with the long fruit axis (upper panel). An inflated cell with identical geometry for the two cases is displayed before growth (left) and after growth (right). For parallel fibers (upper panel), the imaginary fiber length ($^{F}L_1$) is equal before and after growth, as it is orthogonal to the growth direction. Crossed fibers (lower panel) have a non-null projection along the direction of growth, so that the change in width ($b'_c$) exerts a stretch on them, causing an increase in the fiber tension (the tension along the material anisotropy for each layer). This increase in fiber tension is parallel to the fiber direction (the material anisotropy for each of the fiber-dominated layers), and hence has a non-null longitudinal component, which is additional to the tension caused by turgor pressure per se. **(E)** Comparison of growth models using simple transversely isotropic material (left) versus multi-layers (right) in the case of zero Poisson ratio, where the two approaches become almost perfectly comparable (see Supplementary Materials, Star Methods). Deformation and Cauchy stresses are indistinguishable between the two different growth simulations. **(F)** Growth simulated in a block of cells in the multi-layer model (identical to Fig. 4E), but with fibers parallel to the long fruit axis (0º). Initial and final stages of the growth simulation, with ends fixed versus free (stretch ratio 0.86), are shown as in Fig. 4E. Length contraction of the cell block is enhanced with crossed fibers (stretch ratio 0.82, Fig. 4E, 14º with respect to vertical) compared to parallel fibers (stretch ratio 0.86). Single white lines indicate direction of material anisotropy in each simulation (A, E), double white lines indicate direction of fiber orientation in each multi-layer simulation (C, E-F). All heatmaps display trace of Cauchy stress.

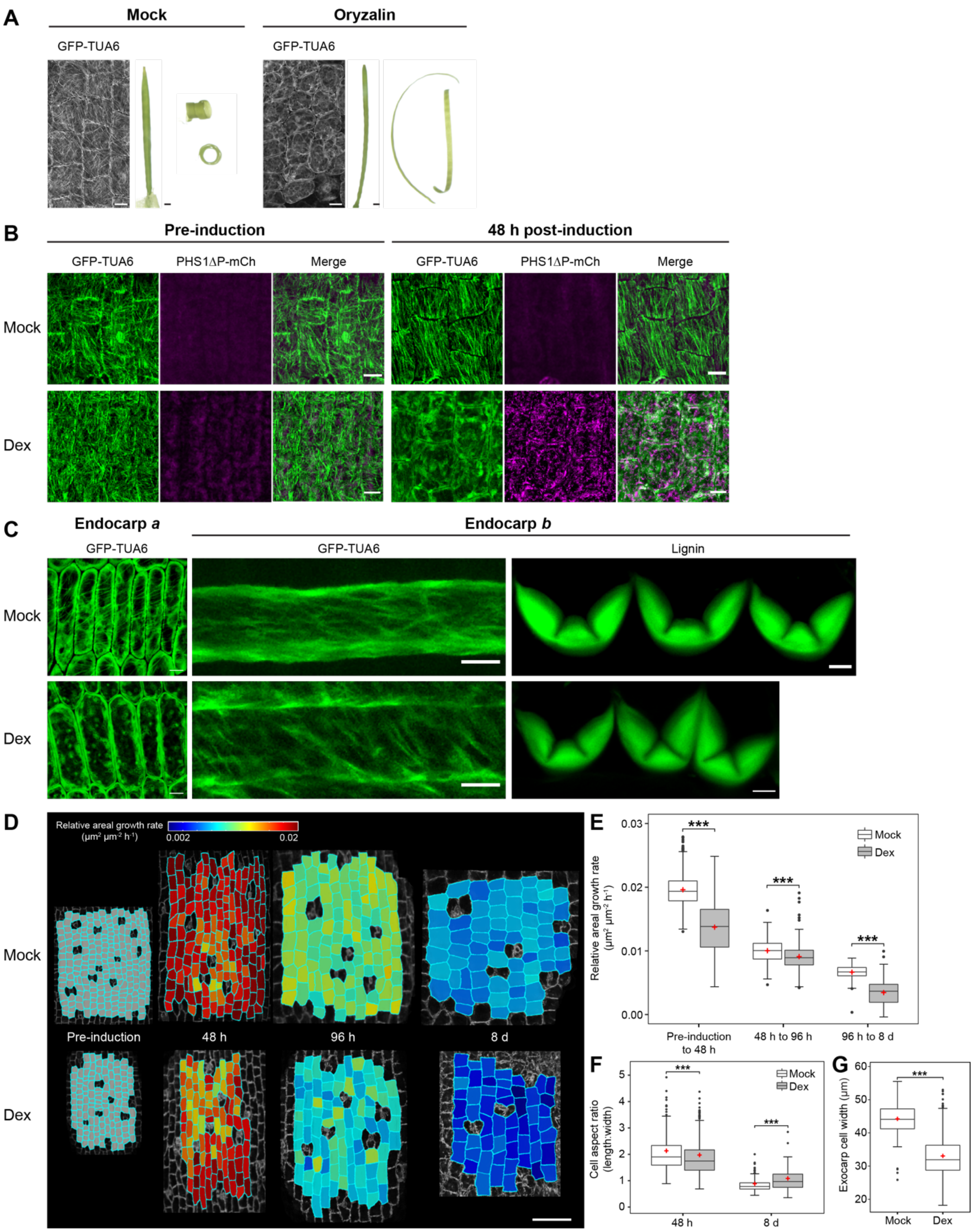


**Figure S3. Exocarp microtubule perturbations, related to Fig. 5. (A)** From left to right, CLSM images of microtubules (GFP-TUA6), whole fruit, and dehisced valves, in mock- vs oryzalin-treated (125 µM Oryzalin, 0.03 % Silwet) *C. hirsuta* fruit. **(B-G)** Live CLSM imaging and quantitative analysis of *C. hirsuta pML1::LhGR>>PHS1ΔP:mCherry; 35S::GFP:TUA6* fruit. **(B)** Exocarp cells of mock- and dexamethasone (Dex)-treated fruit at 0 h (pre-induction) and 48 h post-induction showing microtubules (GFP-TUA6, green) and PHS1ΔP:mCherry (magenta). PHS1ΔP:mCherry expression is only detected in Dex-treated fruit (48 h post-induction), resulting in microtubule depolymerization. **(C)** Endocarp *a* (epidermal layer of the valve endocarp) and endocarp *b* (subepidermal layer of the valve endocarp) cells of mock- and Dex-treated fruit. Microtubules are depolymerized in endocarp *a*, but not endocarp *b* cells in Dex-treated fruit. CLSM images of fixed cross-section of endocarp *b* cells showing the lignified cell wall (stained with basic fuchsin) in mock- and Dex-treated fruit. **(D)** Heat maps of relative areal growth rate of exocarp cells between two consecutive time-points of mock- and Dex-treated fruit. **(E-G)** Boxplots of relative areal growth rate, $n$ = 2226 cells (E), cell aspect ratio (length:width), $n$ = 1571 cells (F), and cell width, $n$ = x cells (G) of exocarp cells in mock- and Dex-treated fruit. Red crosses represent the mean. Significant differences (***) at $p < 0.001$ using a Wilcoxon signed-rank test are indicated. Scale bars: 1 mm (A), 10 µm (B), 5 µm (C-D).

### Supplementary Movies

**Movies S1-S3. Cortical microtubule and CESA3 dynamics in *C. hirsuta* exocarp cells, related to Fig. 3.** Confocal time-lapse series movies of GFP-CESA3 (green) and mCherry-TUA6 (magenta). The smaller green punctae are GFP-CESA3 particles localized at the plasma membrane. Movies are recorded at five frames per second. Acquisition time in seconds (s) is indicated in each movie. Fruits of 7 mm length were selected for the experiment and different fruit were imaged at 0 hours (h) (Movie S1), 48 h later (Movie S2) and 96 h later (Movie S3). Average projections of these movies are shown in Fig. 3E. Scale bars: 5 µm (Movies S1-S3).
