## Supplementary material for "Growth and tension in explosive fruit": STARMethods

### STAR★METHODS

### KEY RESOURCES TABLE

| REAGENT or RESOURCE | SOURCE | IDENTIFIER |
|---|---|---|
| **Experimental Models: Organisms/Strains** | | |
| *A. thaliana*: Col-0 | NASC | |
| *C. hirsuta*: Ox | [1] | |
| *C. hirsuta: pML1::LhGR>>PHS1ΔP:mCherry; 35S::GFP:TUA6* | This study | |
| *C. hirsuta: pCESA3::GFP-CESA3*; *pUbi10::mCherry:TUA6* | This study | |
| **Chemicals** | | |
| Propidium iodide (PI) | Sigma-Aldrich | 25535-16-4 |
| Murashige and Skoog (MS) medium | Duchefa | M0222 |
| Dexamethasone (water soluble) | Sigma-Aldrich | D2915-100MG |
| NaCl | Roth | 231-598-3 |
| Oryzalin | ChemService | N-12729-100MG |
| **Software and Algorithms** | | |
| Fiji | | https://fiji.sc/ |
| MorphoGraphX | | https://www.MorphoGraphX.org |
| MorphoMechanX | | https://www.MorphoGraphX.org/MorphoMechanX |
| Inflation & growth models for MorphoMechanX | This study | https://gitlab.mpcdf.mpg.de/g-gabriella-mosca1/cardaminecontractility_growth.git |
| RStudio | Posit | https://posit.co/ |

### RESOURCE AVAILABILITY

#### Lead Contact

Further information and requests for resources and reagents should be directed to and will be fulfilled by the

**Material Availability**

All materials are available upon request.

**Data and Code Availability**

For the computational simulation, the model source code, the instructions and all templates to reproduce the results can be found here: https://gitlab.mpcdf.mpg.de/g-gabriellamosca1/cardaminecontractility_growth.git

### EXPERIMENTAL MODEL AND SUBJECT DETAILS

**Plant materials and growth conditions**

*Cardamine hirsuta* (Ox), herbarium specimen voucher Hay 1 (OXF) [1], and *Arabidopsis thaliana* (Col-0) were used throughout this study. Plants grown on soil were cultivated in the greenhouse in long-day conditions (days: 20°C, 16 h; nights: 18°C, 8 h). Transgenic plants were generated by the floral dip method using *Agro-bacterium tumefaciens*. *C. hirsuta 35S::GFP:TUA6* plants were described previously [2].

### METHOD DETAILS

**Generation of transgenic lines**

*pML1::GR-LhG4/pOp6::PHS1ΔP:mCherry* was generated as a multiple expression GreenGate cassette from two intermediary modules, built using previously described entry vectors [3-5] (PHS1ΔP, gift from A. Maizel; all other vectors, Addgene). Both modules were combined in pGGZwf01 [6] and transformed into *C. hirsuta 35S::GFP:TUA6* plants [2]. Twenty-eight Basta-resistant lines were selected, tested for dexamethasone induction, and transgene copy number determined (iDNA Genetics). *pUBIQUITIN10::mCherry:TUA6* was generated by *Not*I digest of pMT813 to release the expression cassette. The insert was blunted, ligated into *Sma*I-digested pPZP200-Basta binary vector, and transformed into *C. hirsuta* plants. Transgene copy number was determined (iDNA Genetics) for nine independent lines. A *pCESA3::GFP-CESA3* construct [7] (gift from A. Sampathkumar) was transformed into *C. hirsuta pUbi10::mCherry:TUA6* plants. Ten hyogromycin-resistant lines were selected, analyzed for GFP-CESA3 localization at the plasma membrane, and transgene copy number determined (iDNA Genetics). Representative lines for each construct were analyzed in the T3 generation.

**Microscopy and quantitative image analysis**

A Leica TCS SP8 was used for Confocal Laser Scanning Microscopy (CLSM) with a Nikon 20x water dipping objective (20x/0.95 water) or a Nikon 63x oil objective (63x/1.4 oil) and with the following excitation (ex) and emission (em) parameters (wavelength in nm): GFP ex: 488, em: 550-650 bandpass filter; mCherry and basic fuchsin ex: 561, em: 600-665 bandpass filter; PI ex: 461, em: 550-650 bandpass filter.

**Time-lapse imaging and dexamethasone induction of *pML1::LhGR>>PHS1ΔP:mCherry*:** *C. hirsuta pML1::LhGR>>PHS1ΔP:mCherry; 35S::GFP:TUA6* fruit of 7 mm length were carefully taped to a glass slide, while still attached to the plant, and 0.6 µm z-stack slices of GFP signal in exocarp cells were acquired by CLSM using a 20x/0.95 water dipping lens (0 h time-point, pre-induction). After imaging, plants were separated into treatment and mock groups and fruit were dipped for ten seconds in a solution containing 0.02% Silwett L-77 with 1mM water-soluble dexamethasone (Sigma-Aldrich) (Dex) or without (Mock). Plants were returned to the greenhouse overnight and the same treatments repeated the following day. At the 48 h time-point, CLSM imaging of the same fruits was repeated. After imaging, treatment of the fruits was repeated, and plants were returned to the greenhouse. Treatment of fruits continued every 48 hours and CLSM imaging of the same fruits was repeated at 96 h and 8 d time-points. A minimum of four fruit valve replicates were imaged throughout each time-lapse experiment. Z-stack images were loaded into MorphoGraphX and segmented using signal from the GFP channel [8]. Parent relations for each cell between successive time points were determined and heat maps displayed on the second of two time points. Microtubule orientation, growth anisotropy and all other cellular parameters were quantified using existing functions in MorphoGraphX [8]. The principal direction of growth was determined relative to the fruit long axis.

At the 96 h time-point, GFP-TUA6 signal in the endocarp *a* and *b* cell layers was imaged in detached valves from dex- and mock-treated fruit from the experiment. Whole valves were mounted in water between a glass slide and coverslip and 0.3 µm z-stack slices of GFP signal were acquired by CLSM using a 25x/0.95 water dipping lens.

At the 11 d time-point, dex- and mock-treated fruit from the experiment were triggered to explode. The coiled valves were photographed and circles were fitted to the maximum and minimum curvature of each valve using Fiji [9]. The radius of each circle was measured and valve curvature was quantified as the reciprocal of the radius of curvature.

At the 11 d time-point, detached valves from dex- and mock-treated fruit from the experiment were also embedded in 5% agarose. 100-150 µm sections were cut with a Leica Vibratome VT1000 S, stained for lignin using ClearSee with 0.2% Basic Fuchsin as previously described [6], and 0.5 µm z-stack slices of Basic Fuchsin signal were acquired by CLSM using a 20x/0.95 water dipping lens.

**Osmotic treatments:** Valves were detached from stage 15 (7 mm length) or stage 17b (20-22 mm length) *C.*

*hirsuta 35S::GFP:TUA6* fruit and incubated in 1 mg/mL propidium iodide (PI, Sigma-Aldrich) for ten minutes. Valves were then rinsed in water, cut into smaller sections and adhered with silicone vacuum grease to a small petri dish with the exocarp facing up, and covered with water. The petri dish was moved to the CLSM stage and 0.6 µm z-stack slices of PI signal were acquired, starting at the exocarp cell surface, by CLSM using a 20x/0.95 water dipping lens (high turgor). Water was then removed from the petri dish, replaced with 1 M NaCl and valves were incubated for 60 minutes before imaging of the same cells was repeated (low turgor). Plasmolysis was verified by checking that microtubules were fully depolymerized. Six replicate experiments were performed. Z-stack images were loaded into MorphoGraphX and segmented using signal from the PI channel [8]. Parent relations for each cell between high and low turgor treatments were determined and heat maps displayed on high turgor images. The orientation and magnitude of PDG values were determined relative to the fruit long axis with existing functions in MorphoGraphX [8]. Osmotic experiments using mock- and dexamethasone-treated fruit of *pML1::LhGR>>PHS1ΔP:mCherry; 35S::GFP:TUA6* plants were performed as described above. Four mock- and three dexamethasone-treated replicates were analyzed at 11 d post-induction.

**GFP-CESA3 live imaging and analysis:** Valves were detached from stage 15 (7 mm length), stage 16 (10-12 mm) or early stage 17a (17-19 mm) *C. hirsuta CESA3::GFP-CESA3; Ubi10::mCherry:TUA6* fruit. These fruit stages match the 0 h, 48 h and 96 h timepoints in the timelapse experiment described above. Older fruit did not have GFP-CESA3 signal at the plasma membrane. Valves were cut into smaller sections, mounted in perfluoroperhydrophenanthrene (Sigma-Aldrich) between a glass slide and coverslip, and GFP and mCherry signals were acquired by CLSM using a 63x/1.4 oil immersion lens. A single z-slice was taken every 12 seconds for 30 time-frames. Live imaging was performed for a minimum of 34 cells at each fruit stage. These two-channel images were processed using Fiji [9]. The Stackreg function was used to correct for sample drift and then split into two separate channels. Samples were corrected using the Bleach Correction function followed by background subtraction (50 pixel rolling ball radius). An average projection of the time series was then taken and the brightness/contrast was adjusted. Images were rotated to ensure that the cell long axis is parallel to the fruit long axis. TUA6 and CESA3 trajectory orientations were measured using FibrilTool [10] and the angle between the orientation and fruit long axis was measured.

**Oryzalin treatments:** *C. hirsuta 35S::GFP:TUA6* fruit of 7 mm length were dipped in 125 µM Oryzalin or mock solution (0.03 % Silwet) for 10 min each day for 16 days. GFP-TUA6 was imaged by CLSM, as described above, to verify CMT depolymerization. Whole fruit and dehisced valves were photographed after 16 days.

**Statistical analyses**

The following statistical analyses were done with R Statistical Software: Shapiro-Wilks, Wilcoxon signed-rank, F-test, Student's *t*-test, Pearson's correlation, and graphs were produced using the ggplot2 package [11].

**Computational FEM simulations**

All Finite Element Method simulations were performed within the MorphoMechanX framework (www.MorphoMechanX.org), a modeling platform based on MorphoDynamX and evolved from MorphoGraphX (www.MorphoGraphX.org). Simulation templates were created with the Cell Maker Plugin of MorphoDynamX by using the Block Cell Layers feature and editing a text file to adjust for the different cell templates (see the Git repository at https://gitlab.mpcdf.mpg.de/g-gabriellamosca1/cardaminecontractility_growth for detailed instructions and example files). The reference template consisted of a line of 5 cells connected in the length (y-axis) direction, each cell has the dimensions 26×50×10 µm in width (x-axis), length (y-axis) and depth (z-axis) respectively. All simulations used the aforementioned template, except where cell width or depth was varied (Fig. 2D-E, Fig. S1C), or in the cell block simulations where cells were arranged in a staggered 5×5×5 block of files (Fig. 4E, Fig. S. 2F). Templates were all triangulated with a regular mesh where the elements biggest side is equal to 1 µm. When two cells share a face, this is not duplicated, instead the same mesh elements and nodes refer to both cells. All simulations use a membrane approximation to represent the cell wall, with plane stress and a zero transversal shear strain hypothesis (Kirchhoff-Love theory for membranes, see [12]). Specific material properties assigned are listed in (StarMethods Table 1).

**Inflation simulations (Fig. 2, Fig. S1):** Before the inflation simulation, the ends of the cell file (anticlinal walls orthogonal to the y-axis and belonging to only one cell) are prevented from being able to move in the y-direction (Dirichlet boundary condition). Afterwards, the template is inflated and the Cauchy stress can be visualized. For such a simulation with fixed ends, the maximal Cauchy stress component coincides with the longitudinal stress (parallel to the y-axis in our template orientation), therefore, this can be visualized as well. To know the maximal stress orientation and magnitude of the Cauchy stress tensor, its eigenvectors and values have been computed using the GSL library, specifically the gsl_eigen_symmv function (https://www.gnu.org/software/gsl/doc/html/eigen.html), which works for real, symmetric matrices [13].

Once the inflation process has reached convergence (see the following paragraph for the criterion used), the template ends are allowed to move in all space directions again, and contraction takes place. The stretch ratio is obtained by measuring the length (along y-axis) of the three central cells before inflation, and after the cell ends are released from any constraint (Fig. S1A).

The inflation simulations use a Saint Venant-Kirchhoff transversely isotropic material (the material properties have rotational symmetry around a specific direction, which is the direction conferring anisotropy to the wall, and which can be interpreted as the one reinforced by fibers, even if the description is purely continuous here) and have been developed adopting the same mathematical formulation and computational scheme reported in [2]. There is a small difference for the convergence tolerance, which now requires the mean of average residual force norm and maximal residual force norm to both be below an assigned threshold (StarMethods Table 1). Material anisotropy was assigned on the cell faces, before inflation and growth, as indicated (StarMethods Fig. 1).

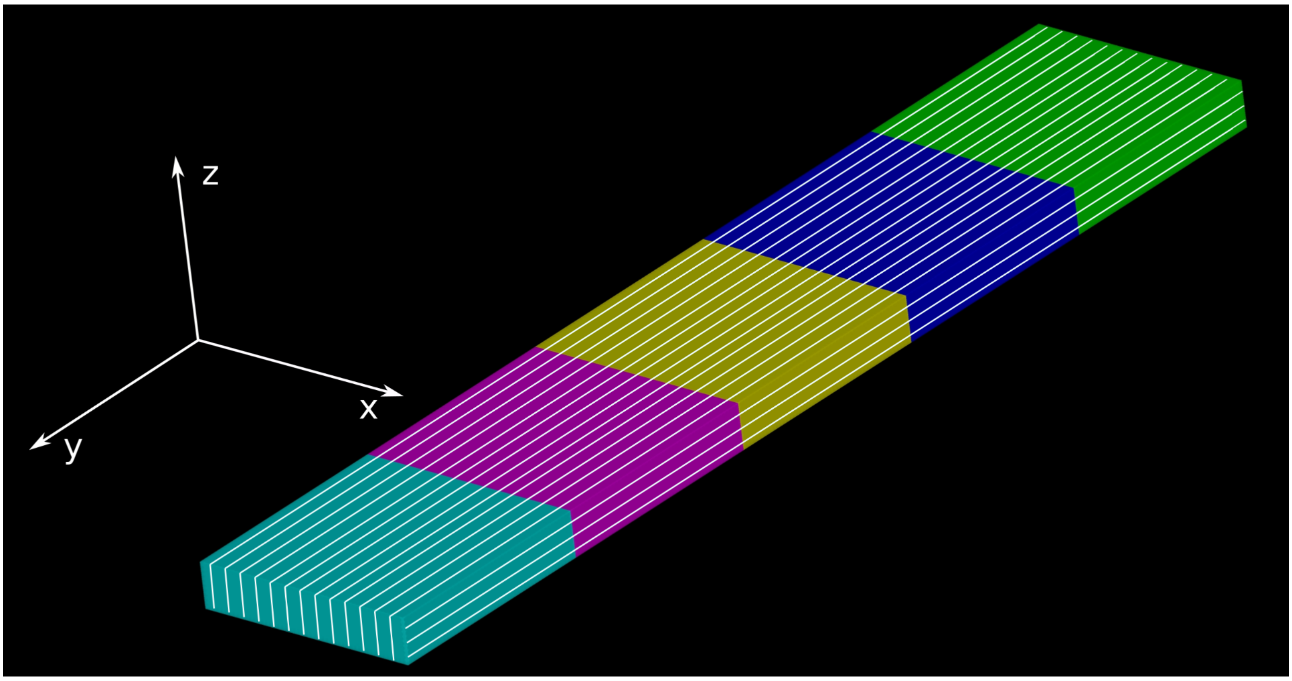


**StarMethods Fig. 1.** A line of cells, as used in the inflation simulations before any deformation occurs, showing how material anisotropy (white lines) is oriented on the different cell walls (parallel to the y-axis on the periclinal walls and the longitudinal anticlinal walls, parallel to the z-axis on the transverse anticlinal walls).

**Single layer growth simulations (Fig. 4A-B, Fig. S2A):** These growth simulations use the same formulation for material description and elastic deformation reported in the previous section. Templates start with a file of cells of sizes 26×50×10 µm in width, length and height respectively. The cell file ends are initially fixed in the y-direction as previously described. After elastic equilibrium has been reached, an iterative growth process takes place (one step of growth, followed by mechanical equilibrium re-computation) until the template reaches a width of approximately 50 µm; during this process the cell file ends are kept fixed in the y-direction. Growth is occurring as a turgor-driven, strain-based relaxation of the reference configuration and acts only as a planar tensor in the element local coordinates (no in-thickness growth). Details about the mathematical formulation and its implementation in MorphoMechanX are provided (Supplementary Information). At the end of the growth phase, the cell-file ends, which were prevented from moving in the y-direction, are released, and the cell file is free to elastically contract.

**Multi-layer growth simulations (Fig. 4C-E, Fig. S2C-F):** Multi-layer growth simulations are conceptually identical to the simulations described in the previous section, with the difference that the material used to describe the cell wall is now comprised of three layers (Fig. S2B). This means that each triangle element of the mesh is assigned three different reference configurations (identical at simulation start), three different material laws and three different growth rules, but share the same current configuration (given by the space coordinates of the triangle vertices, see Supplementary Information Fig. 1).

Since membrane elements have a purely virtual thickness, each layer can be thought of as fully co-penetrating the others and vice-versa, without any notion of spatial hierarchy. While the current configuration is unique and shared with all the layers, MorphoMechanX enables the user to identify each layer in the GUI and assign to it its own reference configuration, material properties and growth rule. In the specific three-layer model used in this work, there are:

(i) Two layers with transversely isotropic Saint-Venant Kirchhoff material properties [2]. For each layer, the anisotropy direction $\mathbf{M}$ (Supplementary Information) was assigned at an angle $\gamma$ with respect to the long axis of the cell file at the simulation start (y-axis in StarMethods Table 1); this angle should not be confused with the angle $\alpha$ made by $\mathbf{M}$ with respect to the first triangle side in the reference configuration (Supplementary Information). For all simulations, but one case (Fig. S2C), one fiber layer was displaced by an angle $+\gamma$, and the other by an angle $-\gamma$, with respect to the y-axis, so that the fibers make a symmetric crossed pattern. The stiff component of each of these layers has a Youngs modulus ($E_{fiber}$) equal to half the value assigned to the Youngs modulus of the stiff component in the single layer growth model. The soft component Youngs modulus ($E_{iso}$) is assigned to be extremely soft (Supplementary Table1), as its value should be negligible with respect to the stiffness of the isotropic layer. These two transversely isotropic layers are not undergoing any growth process, only purely elastic deformation.

(ii) One isotropic layer with Sain-Venant Kirchhoff material properties. The Youngs modulus (E) of this layer has the same value as the soft component ($E_{iso}$) of the single layer growth model. This layer is undergoing turgor-driven, strain-based growth.

All three layers are assigned to be rather incompressible (Poisson ratio $\nu = 0.4$) and turgor pressure acts on one layer (it is irrelevant which one), as its mechanical effect in terms of deformation is then affecting equally all the layers since they share the same unique current configuration.

Computation of the elastic equilibrium for the multi-layer in MorphoMechanX: Given the current configuration (unique) of the mesh triangles, and each layer's reference triangle and material properties, the nodal reaction forces for each layer are computed as usual [2]. In the assembly of the force vector (Voigt notation), the nodal forces from the three layers are summed, the same holds for the "stiffness matrix" used in the semi-implicit Euler scheme [14], whose entries are given by the variation of the nodal forces with respect to a space variation of a mesh node (which means that the contribution of the different layers for the same two nodes involved in the computation is summed). For further details see (Supplementary Information).

Computation of growth for the multi-layer in MorphoMechanX: As already mentioned, only the isotropic layer performs growth, and this is computed as explained above for single layer growth simulations (see also Supplementary Information). This means that only for this layer the reference configuration will change, while for the other two layers it will stay the same from the simulation start until the end.

**StarMethods Table 1: Summary of parameters used in FEM simulations**

| Parameter | Symbol | Value | Main Text Figure/Suppl. Figure |
|---|---|---|---|
| All simulations | | | Fig. 2(C-E), Fig. 4, Fig. S1, Fig. S2 |
| Thickness (single walls) | $t$ | 0.2 $\mu$m | |
| Pressure | P | 0.7 MPa | |
| Mesh element size (longest edge) | | 1 $\mu$m | |
| Convergence tolerance | tol | 1.e-5 | |
| Osmotic treatment simulations | | | Fig. 2(C-E), Fig. 4A, Fig. S1 |
| Young's modulus | $E_{iso}$ | 150 MPa | |
| Young's modulus | $E_{fiber}$ | 4000 MPa | |
| Poisson ratio† | $\nu$ | 0.4 | |
| Cell size | $l \times w \times d$ | (50×10×10) $\mu$m | |
| Cell size | $l \times w \times d$ | (50×25×10) $\mu$m | |
| Cell size | $l \times w \times d$ | (50×40×10) $\mu$m | |
| Cell size | $l \times w \times d$ | (50×50×10) $\mu$m | |
| Cell size | $l \times w \times d$ | (50×50×50) $\mu$m | |
| Growth simulations Single Layer | | | Fig. 4B, Fig. S2A |
| Young's modulus | $E_{iso}$ | 150 MPa | |
| Young's modulus | $E_{fiber}$ | 4000 MPa | |
| Poisson ratio† | $\nu$ | 0.4 | |
| Strain Based Growth Coefficient | $K_{strain}$ | 1 | |
| Strain Growth Threshold (High) | $\epsilon_{thresh}$ | 0.1 | |
| Strain Growth Threshold (Low) | $\epsilon_{thresh}$ | 0.01 | Fig. S1 |
| Growth Step increment | $dt$ | 0.1 | |
| Cell size | $l \times w \times d$ | (50×26×10) $\mu$m | |
| Growth simulations Three Layers | | | Fig. 4(C-E), Fig. S2 (C-F) |
| Cell size | $l \times w \times d$ | (50×26×10) $\mu$m | |
| Isotropic Layer | | | |
| Young's modulus | $E_{iso}$ | 150 MPa | |
| Young's modulus | $E_{fiber}$ | 150 MPa | |
| Poisson ratio† | $\nu$ | 0.4 | |
| Strain Based Growth Coefficient | $K_{strain}$ | 1 | |
| Strain Growth Threshold | $\epsilon_{thresh}$ | 1e-6 | |
| Growth Step increment | $dt$ | 0.5 | |
| Fiber layer (1 & 2) | | | |
| Young's modulus | $E_{iso}$ | 0,1 MPa | |
| Young's modulus | $E_{fiber}$ | 2000 MPa | |
| Poisson ratio† | $\nu$ | 0 | |
| Validation multi-layer with inflation | | | Fig. S2 E |
| Cell size | $l \times w \times d$ | (50×25×10) $\mu$m | |
| Single Layer | | | |
| Young's modulus | $E_{iso}$ | 150 MPa | |
| Young's modulus | $E_{fiber}$ | 4000 MPa | |
| Poisson ratio† | $\nu$ | 0 | |
| Three Layers | | | |
| Isotropic Layer | | | |
| Young's modulus | $E_{iso}$ | 150 MPa | |
| Young's modulus | $E_{fiber}$ | 150 MPa | |
| Poisson ratio† | $\nu$ | 0 | |
| Fiber layer (1 & 2) | | | |
| Young's modulus | $E_{iso}$ | 0.1 MPa | |
| Young's modulus | $E_{fiber}$ | 2000 MPa | |
| Poisson ratio† | $\nu$ | 0 | |

† transversely isotropic material has two independent Poisson ratios, which can be chosen to be $\nu_{fiber-iso}$ (the negative ratio between the deformation induced along the fiber direction and the direct deformation in the isotropic plane, $\nu_{xy}$ in [2]) and $\nu_{iso}$ (the negative ratio between induced and primary deformation both in the isotropic plane, $\nu_z$ in [2]), following the reasoning in [2], those values are assigned so that, at first order, the total cell wall compressibility $\left(\frac{\Delta V}{V}\right)$ for the transversely isotropic material is the same as an isotropic case with Poisson ratio $\nu$. The shear modulus is also assigned following the argument in [2], so that it is no longer an independent parameter.

### SUPPLEMENTARY INFORMATION

Supplementary Figures S1-S3, Supplementary Information, Supplementary Movies S1-S3.
