## Supplementary Information for "Growth and tension in explosive fruit"

### SUPPLEMENTARY INFORMATION OF: GROWTH AND TENSION IN EXPLOSIVE FRUITS

#### CONTENTS


1 Assignation of the material anisotropy vector 2
2 Mathematical formulation of strain-based growth 2
2.1 Implementation of strain-based growth in MorphoMechanX . . . . . . 3
3 Forces and Hessian Assembly in multi-layer 4


#### LIST OF FIGURES


Figure 1 Diagram representig (top left) the current configuration of one mesh triangle as indicated by its global space-coordinates, (top right) the current configuration rotated in the x-y plane by a rotation matrix which alignes the triangle normal to the z-axiz, (bottom) the reference configuration created from the ordered triangle rest lengths ($l_1$, $l_2$, $l_3$), with nodes indicated by $\mathbf{q_1}$, $\mathbf{q_2}$, $\mathbf{q_3}$, and material anisotropy vector $\mathbf{M}$, with angle $\alpha$ w.r.t. the first triangle side $l_1$. . . . . . . . . . . . . . . . . . . . 5

### 1 ASSIGNATION OF THE MATERIAL ANISOTROPY VECTOR

Material anisotropy orientation is assigned by the user through the GUI on the current (or initial) configuration by specifying a vector by means of its x-y-z coordinates w.r.t. the fixed reference frame (see 1). This vector, after normalization, which will be termed $\widetilde{\mathbf{M}}$, is projected on each selected triangle of the mesh. If the projected vector norm is smaller than a prescribed tolerance, there will be no assigned direction on that element and isotropic properties will be prescribed (given by the arithmetic average of the values).

After some manipulation (rotation in the 2D plane of the current configuration triangle by means of a rotation matrix - $M_{3to2D}$ - and creation, from the ordered reference lengths, of the reference triangle, see Sec. 2.1 for details), the 2D deformation gradient of each triangle $\mathbf{F}$, from reference to current (rotated in plane) configuration is computed and, by polar decomposition [Holzapfel, 2002], the pure rotational part of $\mathbf{F}$ is isolated, and this will be named $\mathbf{R}$. The inverse of $\mathbf{R}$ is applied to the anisotropy direction $\widetilde{\mathbf{M}}$ after this as well has been rotated in the plane with $M_{3to2D}$, so that, the anisotropy direction mapped to the reference triangle results:

$$\widetilde{\widetilde{\mathbf{M}}} = R^T(M_{3to2D})\widetilde{\mathbf{M}} \tag{1}$$

Once $\widetilde{\widetilde{\mathbf{M}}}$ has been normalized, the angle made by it with the first reference triangle side can be computed and stored as material anisotropy attribute of the triangle. The normalized $\widetilde{\widetilde{\mathbf{M}}}$ can be identified with $\mathbf{M}$ of the following Section.

### 2 MATHEMATICAL FORMULATION OF STRAIN-BASED GROWTH

Growth is modeled in an incremental way [Bozorg et al., 2016], where each growth iteration constitutes an update of the resting configuration for an infinitesimal time step $\delta t$ after which the elastic mechanical equilibrium needs to be recomputed to ensure continuity of the current configuration [Rodriguez et al., 1994]. We assume that biologically the time scales for growth are much smaller than those of elasticity [Goriely and Amar, 2007] and this justifies our incremental approach to modeling growth. Following the formalism of Bozorg et al. [2016], the incremental growth tensor $\mathbf{f_g}$ acts on the reference configuration at time t ($\mathbf{X}_0(t)$, so that the update of the reference configuration results:

$$\mathbf{X}_0(t+\delta t) = \mathbf{F_g}(t+\delta t)\mathbf{X}_0(0) = \mathbf{X}_0(t) + \delta t\mathbf{f_g}(t)\mathbf{X}_0(t) \tag{2}$$

Where $\mathbf{F_G}(t)$ is the integrated growth tensor which acts on the original (at $t = 0$) reference configuration $\mathbf{X}_0(0)$ and is in relation to the infinitesimal growth tensor through the following expression, which highlights the exponential nature of growth through a finite amount of time:

$$\mathbf{F_g}(t) = \exp[\int_0^t \mathbf{f_g}(t')dt'] \tag{3}$$

In this context, by "reference configuration" it is not meant directly the body, represented as a (piece-wise) continuous distribution of matter (material points) in space and time, in the absence of external and internal forces, but rather the collection of the tangent spaces (the tangent bundle) at each point $\mathbf{T_P}(t)$ of the embedding in $\mathbf{R}^3$ of the material points [Holzapfel, 2002, Goriely and Moulton, 2011]. In fact the growth tensor, similarly to the deformation gradient, is a mapping (push forward) defined on tangent vectors, in this case from a reference configuration at time t ($\mathbf{X}_0(t)$, to a reference configuration at time $t+\delta t$ ($\mathbf{X}_0(t+\delta t)$, and this tells us how the metric is affected by growth [Vetter et al., 2013].

#### 2.1 Implementation of strain-based growth in MorphoMechanX

In the simulation framework adopted in this work, growth is applied directly to the discretized mesh elements which represent the reference configuration. The mesh is constituted by linear triangular elements, so that the tangent space on each triangle is isomorphic to $\mathbf{R}^2$ (growth orthogonal to the element plane is not considered), hence it is possible to apply the incremental growth tensor $\mathbf{f_g}(t)$ directly to two reference triangle edges (if $\mathbf{q_1}$, $\mathbf{q_2}$, $\mathbf{q_3}$ are the triangle ordered vertices, the edges can be chosen to be $\mathbf{q_2} - \mathbf{q_1}$ and $\mathbf{q_3} - \mathbf{q_1}$) to know the updated reference configuration of the triangle in local coordinates (which are the only ones required for the calculations described below), provided there is a way to store locally information like material anisotropy orientation.

In practice this is achieved by computing a incremental growth tensor which is proportional to the strain in the element (Biot strain tensor, $\mathbf{E_B}$, chosen because it is linear w.r.t. the stretch) and acts on the reference configuration of the mesh triangles.

- For each triangle of the template mesh, the current configuration is stored in terms of nodal coordinates (3D positions) in a fix external Cartesian coordinate frame. The reference element is stored as the assigned reference virtual thickness (which does not change), the ordered rest lengths of the triangle, together with the angle made between the material anisotropy direction and the first side of the triangle, $\mathbf{q_2} - \mathbf{q_1}$ (the triangle is consistently build with a counterclockwise orientation, see Fig.1).

- The reference triangle at a certain time $t$ is reconstructed from its stored reference lengths in a 2D plane ($x - y$ plane): since its position in the plane is irrelevant for the calculations performed, by convention the first side is aligned with the $x$—axis, so that, being $\mathbf{q_1}$, $\mathbf{q_2}$, $\mathbf{q_3}$ the ordered positions of the reference triangle,

$$\begin{aligned} \mathbf{q_1} &= (0, 0) \\ \mathbf{q_2} &= (q_{2x}, 0) \\ \mathbf{q_3} &= (q_{3x}, q_{3y}), \\ \text{with } & q_{2x},\ q_{3y} > 0 \end{aligned}$$

- The current triangle configuration at time $t$, given by its ordered vertices in 3D $\mathbf{p_1}$, $\mathbf{p_2}$, $\mathbf{p_3}$, after a translation which maps $\mathbf{p_1}$ into the origin, is rotated in the $x - y$ plane by computing the triangle normal $\mathbf{n} = (\mathbf{p_2}) \times (\mathbf{p_3})$ and finding the 3D rotation matrix ($M_{3to2D}$) which rotates the normalized vector $\tilde{\mathbf{n}}$ into the unit normal vector $\tilde{\mathbf{z}} = (0, 0, 1)$ (see also Fig. 1).

- The stretch tensor squared ($\mathbf{U}^2 = \mathbf{F}^T\mathbf{F}$, where $\mathbf{F}$ is the purely elastic deformation gradient) is computed using as input the corresponding positions of the reference ($\mathbf{q_1}$, $\mathbf{q_2}$, $\mathbf{q_3}$) and current ($\mathbf{p_1}$, $\mathbf{p_2}$, $\mathbf{p_3}$) triangles, and the standard FEM technique for linear elements in 2D [Zinkiewicz et al., 2005].

- The stretch tensor squared ($\mathbf{U}^2$), which is a 2D symmetric tensor, is easily diagonalized and since it shares the same eigenvectors of the Biot strain tensor ($\mathbf{E_B}$), those are automatically provided, while the eigenvalues of $\mathbf{E_B}$, $\epsilon_{B_i}$, are computed from the eigenvalues of $\mathbf{U}^2$, $u_i^2$:

$$\epsilon_{B_i} = \sqrt{{u_i}^2} - 1$$

- The strain based incremental growth tensor is then built in its diagonal representation:

$$\mathbf{f_g}_D = K \begin{pmatrix} \max(0, \epsilon_{B1} - \epsilon_{thresh}) & 0 \\ 0 & \max(0, \epsilon_{B2} - \epsilon_{thresh}) \end{pmatrix} \quad (4)$$

K is a proportionality coefficient which in all our simulations has been kept uniformly equal to 1, $\epsilon_{thresh}$ is a value that the Biot strain principal values $\epsilon_{B_i}$ need to exceed so that growth acts in the direction of their eigenvector (the operator prevents negative growth from happening). When a higher threshold is set, more stretch is required before growth can occur. In the case of a transversly isotropic material assigned to the wall of a pressurized cell, it is possible to fine tune this parameter so that strain-based growth occurs only in the soft direction.

- To apply the growth tensor to the vectors defining the triangle ($\mathbf{q_2}$ and $\mathbf{q_3}$, since $\mathbf{q_1}$ is set to be the origin), the strain based incremental growth tensor needs to be represented in the same coordinate basis of the reference triangle. This is done using the spectral theorem for symmetric real matrices, so that

$$\mathbf{f_g} = \mathbf{V}(\mathbf{f_g}_D)\mathbf{V}^T \tag{5}$$

where $\mathbf{V}$ is the unitary matrix, whose columns are made by the eigenvectors of $\mathbf{U}$ (or equally $\mathbf{U}^2$ or $\mathbf{f_g}$, as they are the same).

- Finally the update of the reference configuration is computed as:

$$\mathbf{X}_0(t+\delta t) = (\mathbb{1} + \delta t \mathbf{f_g})\mathbf{X}_0(t) \tag{6}$$

where here $\mathbf{X}_0 = (\mathbf{q2}, \mathbf{q3})$ and from the updated reference coordinates, the updated (ordered) reference lengths of the triangles can be recomputed and will be stored.

- The material anisotropy direction, which is stored with the reference configuration, needs to be updated as well. The assumption made is that in the strain-based growth modeled, the anisotropy direction is advected together with the element growth (it might be different if growth occurs by pure accretion). This means that, if $\mathbf{M}$ is the normalized vector of the anisotropy direction (obtained from the angle $\alpha$ made by it w.r.t. the first triangle side, see Fig. 1), it will be so updated:

$$\mathbf{M}(t+\delta t) = (\mathbb{1} + \delta t \mathbf{f_g})\mathbf{M}(t) \tag{7}$$

the vector length is irrelevant as the only information stored will be the updated angle made with the first triangle side.

#### 3 FORCES AND HESSIAN ASSEMBLY IN MULTI-LAYER

As with the simple transversly isotropic material, the equilibrium configuration for the multi-layer is found via a semi-implicit Euler scheme Press et al. [2002], which is iterated over several time steps $t_k$ until convergence has been reached:

$$\hat{u}_{\xi i}(t_{k+1}) = \hat{u}_{\xi i}(t_k) + dt\left(\mathbb{1} - dt\mathbb{H}(t_k)_{\xi i, \theta j}\right)^{-1} \frac{\partial \Pi}{\partial \hat{u}_{\theta j}}|_{t_k} \tag{8}$$

where $\hat{u}_{\xi i}$ is the nodal displacement at node $\xi$, in the space coordinate $i$, $dt$ is the time increment, $\mathbb{H}(t_k)_{\xi i, \theta j}$ is the Hessian of the total potential energy (second derivative of $\Pi$, the total potential energy, made of strain energy and pressure energy potential contribution, w.r.t displacement function) made w.r.t to the displacements at node $\xi$ and $\theta$, space coordinates $i$ and $j$ respectively. As last, $\frac{\partial \Pi}{\partial \hat{u}_{\theta j}}$ is the derivative of the total potential energy w.r.t. nodal displacement. The total potential energy is now contributed in an additive way by the strain energy function of the

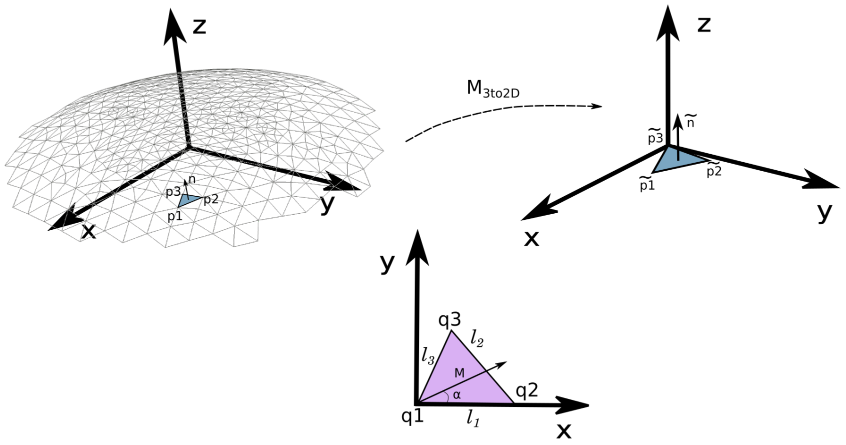


**Figure 1:** Diagram representig (top left) the current configuration of one mesh triangle as indicated by its global space-coordinates, (top right) the current configuration rotated in the x-y plane by a rotation matrix which alignes the triangle normal to the z-axiz, (bottom) the reference configuration created from the ordered triangle rest lengths ($l_1$, $l_2$, $l_3$), with nodes indicated by $\mathbf{q_1}$, $\mathbf{q_2}$, $\mathbf{q_3}$, and material anisotropy vector $\mathbf{M}$, with angle $\alpha$ w.r.t. the first triangle side $l_1$.

three layers (which in the membrane approach are perfectly overlapping), plus the turgor pressure term :

$$\Pi = \Pi_{Iso} + \Pi_{Fiber1} + \Pi_{Fiber2} - \int_{\Omega_0} PJd_0V \tag{9}$$

(with $\Omega_0$ the volume enclosed by the membrane in the undeformed configuration, P the turgor pressure and J the Jacobian of the deformation gradient), it is possible to write derivative of the total potential energy and of the Hessian as a sum of the derivatives (first or second order) of the single layer terms, plus the pressure term. As example one component of the "force vector" $\frac{\partial \Pi}{\partial \hat{u}_{\theta j}}$ is explicitly written:

$$\frac{\partial \Pi}{\partial \hat{u}_{\theta j}} = \frac{\partial \Pi_{Iso}}{\partial \hat{u}_{\theta j}} + \frac{\partial \Pi_{FIber1}}{\partial \hat{u}_{\theta j}} + \frac{\partial \Pi_{Fiber2}}{\partial \hat{u}_{\theta j}} + \frac{1}{3} P \sum_{\theta - neigh} \big(Area(t)\mathbf{n}_j\big) \tag{10}$$

with $\sum_{\theta - neigh}$ indicating the sum over the elements sharing the node $\theta$ of the elemental area A, with normal vector $\mathbf{n}$ and component j, while the Hessian is computed numerically from the force vector just described.

To be able to build Eq. 10, which is exclusively node-based, it is first necessary to compute, element-wise, the elemental nodal force $\frac{\partial \Pi}{\partial \hat{u}}$, which is stored paired to its own element type (Triangle Iso element, Triangle Fiber1 element, Triangle Fiber2 element), making it straightforward to compute then the layer contribution of the force vector to each node, as everything is additive. The pressure term is computed only one and associated arbitrarily to one of the layers.

---
